# Dopamine Release from the Locus Coeruleus Enables Temporal Binding in the Hippocampus

**DOI:** 10.64898/2026.10.13.758657

**Authors:** Ana Paula Crestani, Cassiano Ricardo Alves Faria Diniz, Deger Carda, Savannah Maw, Brian Joseph Wiltgen

## Abstract

The hippocampus forms memories by associating events that are separated in time. These associations can be studied using trace conditioning, a procedure where a temporal gap is inserted between a conditional stimulus and an unconditional stimulus. Here, we examined how dopaminergic projections to CA1 support trace fear conditioning. Using fiber photometry and genetic lesions, we demonstrate that locus coeruleus (LC)-derived dopamine, rather than VTA input, is the primary source of dopamine in CA1 and is essential for trace learning. LC lesions reduced CA1 principal neuron activity and c-Fos expression. In-vivo imaging revealed that successful learning requires post-shock reactivation of stimulus-specific ensembles, a process compromised by LC lesions or long trace intervals. Finally, optogenetic stimulation of LC-CA1 projections during the post-shock period was sufficient to rescue learning in long-trace protocols. These findings identify LC-CA1 dopamine as a critical regulator of temporal binding via the facilitation of hippocampal memory trace reactivation.

## INTRODUCTION

The ability to associate events separated in time is a fundamental cognitive process that enhances survival. In the laboratory, these associations can be studied using trace fear conditioning (TFC), where a neutral conditioned stimulus (CS) is followed by an unconditioned stimulus (US) after a brief delay ^1–4^. Unlike delay conditioning, where the CS and US overlap, TFC requires the brain to maintain a representation of the CS across a temporal gap to form a robust association. While it is well-established that the CA1 subregion of the dorsal hippocampus (dCA1) is essential for bridging these discontinuous stimuli, the precise neuromodulatory mechanisms that sustain this temporal association remain a subject of active investigation ^5,6^.

Temporal constraints are a defining feature of TFC. Research has shown that while a 15- to 30-second gap typically produces robust learning in rodents, the association begins to decay as the trace interval is extended ^7^. Previous studies in mice have demonstrated strong learning at 20-second intervals, but the specific point at which associative learning fails, and the biological factors determining this limit, are not fully understood. Our data confirm that extending this interval to 90 or 120 seconds significantly impairs the formation of associative memories, leading to a marked decrease in conditioned freezing behavior compared to shorter intervals of 20 or 60 seconds. Understanding how the brain overcomes these temporal gaps is critical for elucidating the boundaries of hippocampal-dependent memory.

A leading candidate for modulating hippocampal activity during temporal processing is the dopaminergic system. While the ventral tegmental area (VTA) is the classical source of dopamine (DA) in the brain, recent evidence suggests that the locus coeruleus (LC) serves as a major source of DA in the dorsal hippocampus (dHC)^8–11^. We have previously demonstrated that DA release from the LC to the dCA1 facilitates TFC learning^12^. However, the relative contributions of LC versus VTA projections to the CA1 during the various phases of trace conditioning, and how these projections influence the physiological temporal gap between the tone and the shock, remain unclear.

Beyond simple neurotransmitter release, the efficacy of TFC may depend on the post-shock reactivation of specific neuronal ensembles. Indeed, we have found that CA1 pyramidal neurons responding to the tone or trace interval often show increased activity following the delivery of the US. This post-shock reactivation is thought to stabilize the memory trace, effectively “tagging” the preceding stimuli for associative strengthening ^13^. A similar process takes place during spatial learning, where place cells are reactivated after reward delivery to enhance memory for the path leading to food ^14–17^. It is possible that the failure to learn during long trace intervals or following neuromodulatory disruption is due to a reduction in this reactivation process.

In the current study, we investigated the mechanisms by which LC-derived DA modulates dCA1 activity to promote TFC. Using a combination of fiber photometry, in vivo calcium imaging with miniscopes, and optogenetics, we explored how DA levels and neuronal activity patterns change as the trace interval is prolonged. We hypothesized that dopamine release from the LC, rather than the VTA, is necessary to drive the post-shock reactivation of CA1 pyramidal neurons that is required for successful TFC ^12^.

We first established the temporal limits of TFC in mice, confirming that 120-second intervals impair associative learning. We then demonstrated that selective lesions of the LC-dCA1 projection, but not the VTA, significantly reduce DA release in the hippocampus and impair trace fear learning. Our calcium imaging revealed that while LC lesions do not alter the activity of CA1 interneurons, they significantly reduce the global activity and c-Fos expression of CA1 pyramidal neurons. Crucially, we found that both LC lesions and long trace intervals compromise the post-shock reactivation of tone- and trace-responsive neurons. Finally, we show that optogenetic stimulation of LC-dCA1 projections during the post-shock period is sufficient to rescue associative learning even at 120-second trace intervals. Together, these findings identify the LC-dCA1 dopaminergic pathway as a critical regulator of the post-stimulus neuronal dynamics necessary for bridging events across time.

## RESULTS

### LC lesion, but not VTA lesion, decreases dopamine release in CA1 and impairs trace fear learning

Previous studies suggest that the CA1 subregion of the dHC is required to associate discontiguous stimuli, as in the case of trace fear conditioning ^13,20,21^. In addition, we have recently demonstrated that dopamine release from the LC to the dCA1 facilitates TFC ^12^.

Here, we investigated in detail whether the dopaminergic projection from the LC to dCA1 is indeed essential for TFC. Initially, we used viral vector (flex-taCasp3) injected into the LC to induce Cre recombinase (Cre)-dependent caspase-3 expression in tyrosine hydroxylase (Th)-Cre+ mice. When brought together, these set of tools induce the death of neurons that express Th. An independent group of mice were infused with the same virus into the ventral tegmental area (VTA) in order to compare dopaminergic lesions on dopamine release in dCA1. Neuronal death of transfected neurons was confirmed by *post-mortem* reduction in Th immunostaining (**Fig. 1D-F**). LC and VTA sham surgeries (no viral infusion) were used as a lesion-free control group. All mice received an injection with viral vector (hSyn-DA3h) into the dCA1 to express the dopamine sensor (DA3h) and, immediately afterwards, had a fiber implanted right above the CA1, enabling dynamic measurement of CA1 dopamine release (fiber photometry) throughout the TFC task. **Fig. 1A** describes the experimental design in detail.

**Figure 1.**
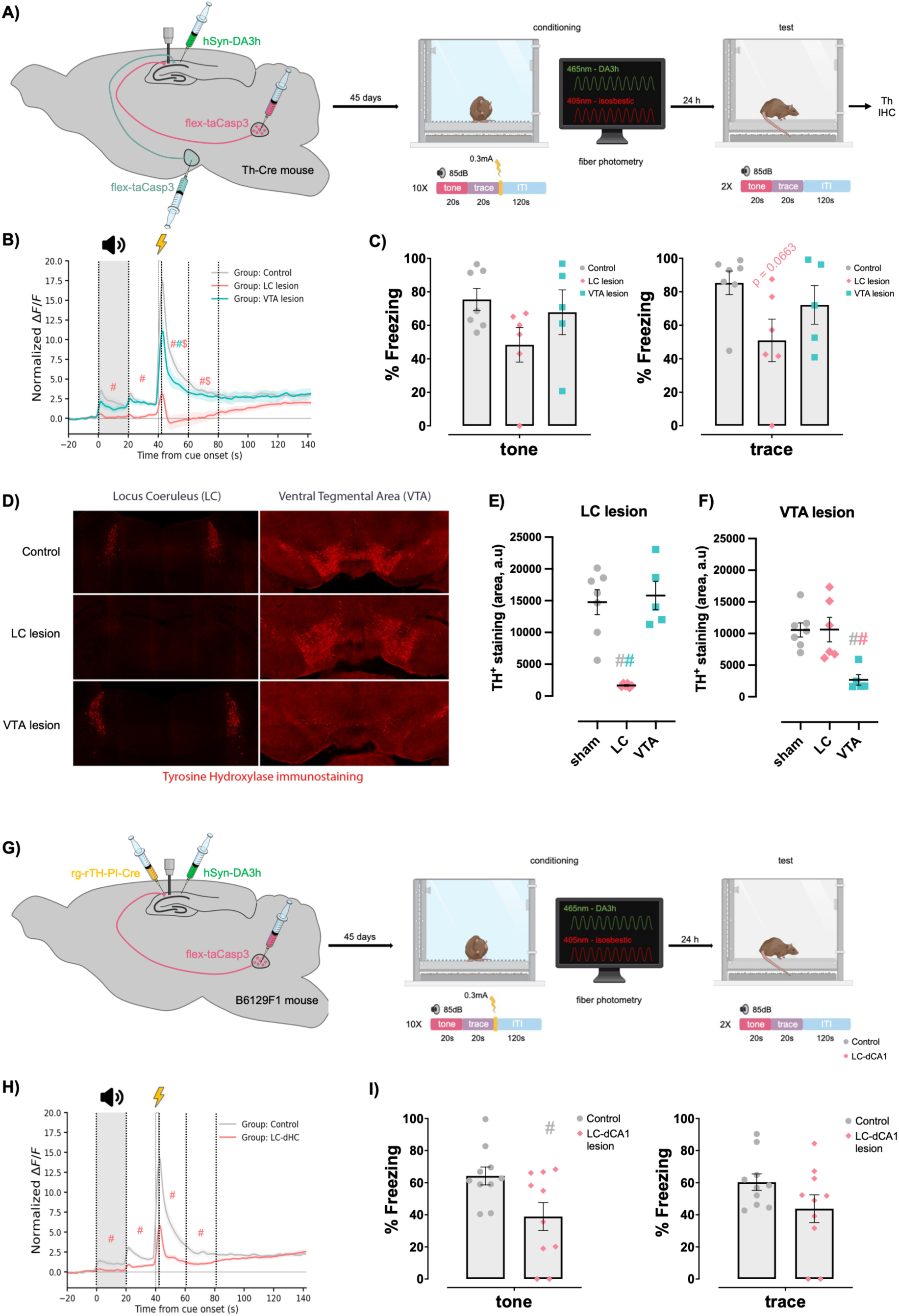
LC lesion, but not VTA lesion, eliminate DA in CA1, impairing trace fear learning. **A)** Dopamine sensor (DA3h) was injected in the dCA1. Cre-dependent caspase-3 was used to lesion the LC (n = 6) or VTA (n = 5) in different groups of Th-Cre mice. Sham animals were used as controls (n = 7). **B)** Fiber photometry traces showing DA release in CA1 throughout TFC. The average of the DA signal was calculated every 20 seconds, corresponding to the baseline, tone, trace, and five post-shock periods. Two-way repeated measures ANOVA showed main effects for Group (F (2,15) = 9.330, p = 0.0023) and Trial Phase (F (7,105) = 37.45, p < 0.0001), as well as an interaction effect between Group x Trial Phase (F (14,105) = 12.10, p < 0.0001). Tukey’s post-hoc test indicated that the DA signal significantly decreased in the LC lesion group during tone, trace, post-shock- 1, and post-shock-2 periods compared to the sham group (represented by salmon #), and during post-shock-1 and post-shock-2 in comparison to the VTA lesion group (salmon $). DA levels were also lower in the VTA lesion group compared to the sham group throughout the post-shock-1 period (cyan #). Furthermore, the sham and VTA groups showed higher DA signals in all trial phases compared to baseline, while the LC lesion group exhibited higher DA levels during post-shock-4 and post-shock-5 compared to baseline. The color of the # symbol indicates which group (salmon: LC lesion and cyan: VTA lesion) had a p-value < 0.05 compared to the sham group. Salmon $ indicates that LC lesion group differed from VTA lesion group (p < 0.05). See also Figure S5. **C)** Freezing during the test session. One-way ANOVA indicated a trend effect during the trace-like period (F (2,15) = 3.013, p = 0.0794). See also Figure S6. **D)** Representative images of tyrosine hydroxylase immunostaining. The images on the left show the LC, while those on the right show the VTA. The Cre-dependent caspase virus visually reduced the expression of tyrosine hydroxylase in the LC (middle left) and VTA (bottom right), respectively. **E)** Quantification of the Th staining in the LC. Linear mixed-effects model revealed a significant effect of LC lesion (z = −5.64, p < 0.001). Control animals (n = 6 animals, 18 observations) showed a mean fluorescence intensity of 14,755.86 ± 1,578.13 (Coefficient ± SE) arbitrary units (a.u.), while LC-lesioned animals (n = 6 animals, 18 observations) exhibited a reduction (β = −13,100.02 a.u., SE = 2,322.94) compared to controls, representing an 89% decrease relative to the control mean. The model also indicated no significant effect of VTA lesion on LC fluorescence intensity (β = 1,033.88, SE = 2,444.83, z = 0.42, p = 0.672). Data is presented as mean fluorescence intensity for each animal. The symbol # indicates p < 0.05, and the color specifies from which groups the LC lesion group differs from. **F)** Quantification of the Th staining in the VTA. Linear mixed-effects model revealed a significant effect of VTA lesion (z = −3.90, p < 0.001). Control animals (n = 6 animals, 18 observations) showed a mean fluorescence intensity of 10,563.19 ± 1,307.16, while VTA-lesioned animals (n = 6 animals, 18 observations) exhibited a reduction (β = −7,900.72, SE = 2,025.04) compared to controls, representing a 75% decrease relative to the control mean. The model indicated no significant effect of LC lesion on VTA fluorescence intensity (β = 58.98, SE = 1,924.09, z = 0.03, p = 0.976). Data is presented as average fluorescence intensity for each animal. The symbol # indicates p < 0.05, and the color specifies which groups the VTA lesion group differs from. **G)** Dopamine sensor (DA3h) and retrograde virus to express Cre in the catecholaminergic neurons (rTH-PI-Cre) were injected in the dCA1. Cre-dependent caspase was infused in the LC to lesion LC-dCA1 projecting neurons. **H)** Fiber photometry traces showing DA3h responses to TFC. The average of the DA signal was calculated every 20 seconds, corresponding to the baseline, tone, trace, and five post-shock periods. Two-way repeated measures ANOVA showed main effects for Group (F (1,17) = 16.31, p = 0.0009) and Trial Phase (F (7,119) = 47.76, p < 0.0001), as well as an interaction effect between Group x Trial Phase (F (7,119) = 14.19, p < 0.0001). Tukey’s post-hoc test indicated that the DA signal significantly decreases in the LC-dCA1 projection lesion group (n = 9) compared to the sham group (n = 10) during tone, trace, post-shock-1, and post-shock-2. # indicates p < 0.05. See also Figure S7. **I)** Freezing during the test session. The LC-dCA1 lesion group (n = 10) showed less freezing compared to the sham group (n = 10) during tone presentation (unpaired t-test, t (18) = 2.450, p = 0.0248). # indicates p < 0.05. See also Figure S8. All data are expressed as mean ± SEM.

The average DA signal was calculated every 20 seconds, encompassing the baseline, tone, trace, and five post-shock periods (post-shock-1, -2, -3, -4 and -5). First, the control and VTA groups depicted an increase in DA signals throughout all trial phases when compared to baseline (**Fig. 1B**). Meanwhile, the LC lesion completely abolished DA release in CA1 during the tone, trace, shock, post-shock-1, and post-shock-2 as evidenced by the signal profile of the control group (**Fig. 1B**). Only after post-shock-2 did the DA signal in the LC lesion group appear to slowly return to normal levels. Conversely, VTA lesion induced a partial decrease in CA1 dopamine release only during shock and post-schock-1, compared to the control group (**Fig. 1B**). Confirming the difference in the effects of LC and VTA lesions on CA1 DA release, the signal from the LC group differs significantly from the VTA group in all trial phases, except from post-shock-3 onwards.

Even though the groups did not differ in freezing levels during the conditioning session (**Fig. S1**) or during the 2 min baseline period of the test session (**Fig. S2**), the group with LC lesion showed a consistent decrease in freezing levels during both the tone period and the trace-like period of the test session (**Fig. 1C**). All lesion groups presented virtually complete neuronal lesion limited to the target region that received flex-taCasp3 viral vector (**Fig. 1D**). The magnitude of neuronal lesion was measured by the reduction in the fluorescence intensity of Th staining. Measurements were performed on 3 brain slices containing the locus coeruleus and 3 slices containing ventral tegmental area for each animal. A linear mixed-effect model was used to account for repeated Th measurements within subjects. Fluorescence intensity in the LC and VTA were considered the dependent variables, with animal was included as a random intercept. The model revealed that fluorescence intensity in LC was significantly reduced by LC lesion (compared to the reference/control group), but not by VTA lesion (**Fig. 1E**). Similarly, VTA lesion decreased fluorescence intensity in the VTA, but not in the LC (**Fig. 1F**).

These results suggest that LC lesions have a much greater influence than VTA lesions on DA released in CA1 during TFC learning and that DA released by the LC is important for such learning to be properly established.

Given that the LC projects to numerous other brain regions besides the hippocampus, and that this matter might be a confounding factor for our results, we subsequently carried out a more refined experimental designed to lesion only LC neurons projecting to the dHC CA1. Thus, a retrograde virus to express Cre in catecholaminergic neurons (rg-rTH-PI-Cre) was injected into the dHC CA1, while the viral vector for Cre-dependent caspase 3 expression (flex-taCasp3) was infused into the LC. Again, all mice received an injection with viral vector (hSyn-DA3h) into the dHC CA1 to express the dopamine sensor (DA3h) and, immediately afterwards, had a fiber implanted right above the CA1. **Fig. 1G** contains details on the experimental design.

In agreement with our previous results, this experiment showed that specific lesioning of CA1-projeting LC neurons reduced DA release in CA1 during tone, trace, shock, post-shock-1, and post-shock-2 (**Fig. 1H**). As expected, the lesioned mice showed significantly lower freezing levels during the test session (**Fig. 2I**), even though freezing levels did not differ between groups during the conditioning session (**Fig. S3**) nor during the 2 min baseline of the test session (**Fig. S4**).

**Figure 2.**
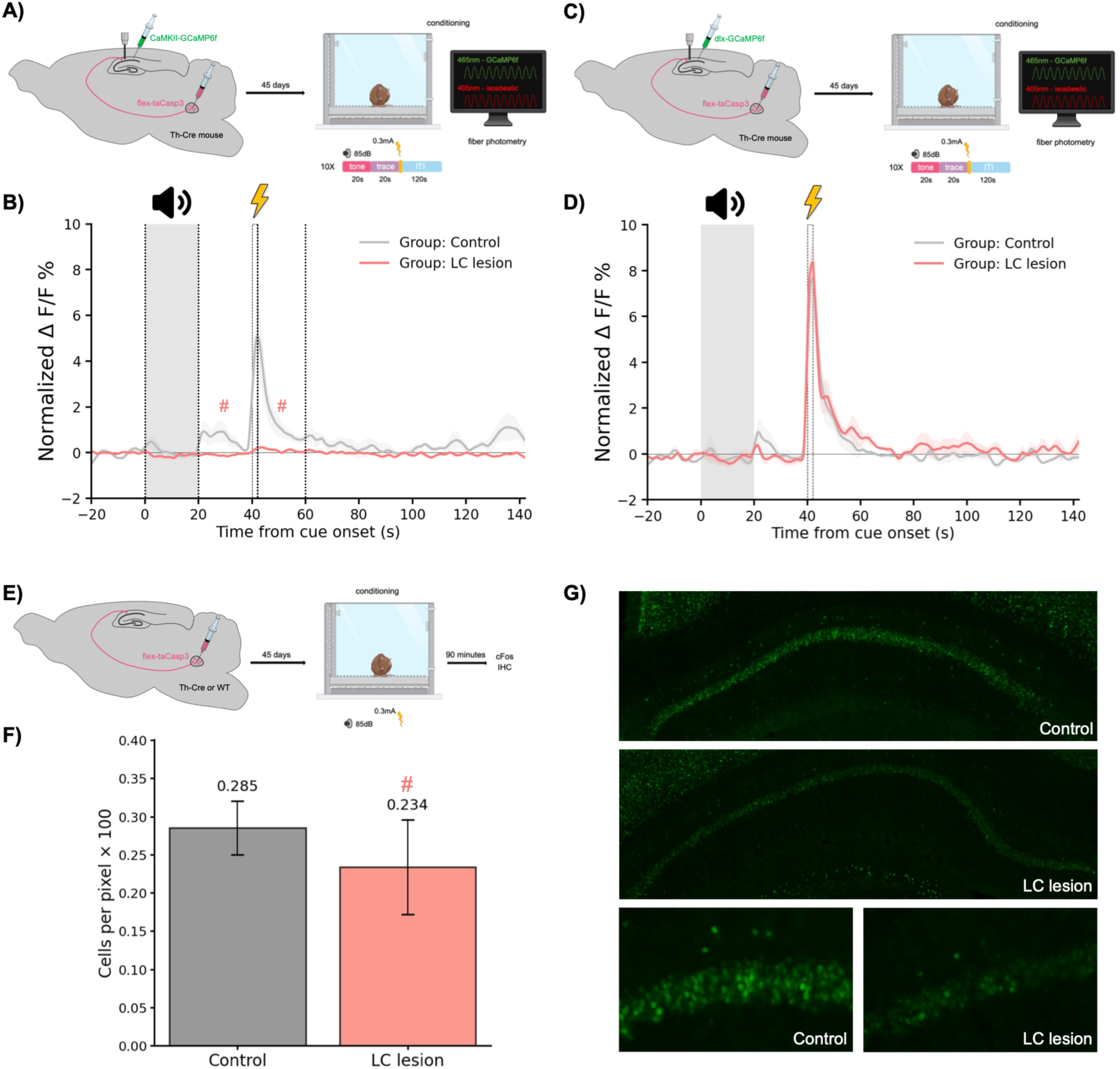
LC lesion reduces c-Fos expression and activity of pyramidal neurons in CA1, while interneurons’ activity remains unaffected. **A)** Cre-dependent caspase was injected into the LC of Th-Cre (LC lesion, n = 3) and WT (control, n = 4) mice, while a fluorescent calcium sensor was expressed in dCA1 pyramidal neurons (CaMKII-GCaMP6f). Following a 45-day gap, bulk calcium activity was measured during TFC using fiber photometry. **B)** Bulk calcium activity of dCA1 pyramidal neurons (CaMKII-GCaMP6f) is reduced during trace and post-shock-1 periods in LC lesion group compared to control mice (Two-way repeated measures ANOVA, main effect of Trial Phase (F (7,35) = 4.726, p < 0.0008) and interaction Group x Trial Phase (F (7,35) = 3.181, p = 0.0103); Tukey’s post-hoc test p = 0.062 and p < 0.0001, respectively). # indicates statistical difference. **C)** Cre-dependent caspase was injected into the LC of Th-Cre (LC lesion, n = 3) and WT (control, n = 3) mice, while a fluorescent calcium sensor was expressed in dCA1 interneurons (dlx- GCaMP6f). Following a 45-day gap, bulk calcium activity was measured during TFC using fiber photometry. **D)** Bulk calcium activity of dCA1 interneurons (dIx-GCaMP6f) is not altered by LC lesion. **E)** The LC of Th-Cre (LC lesion) and WT (control) mice were injected with a Cre-dependent caspase. Following a 45-day gap, mice underwent TFC and were perfused 90 minutes later. **F)** Quantification of cFos immunostaining in CA1. Linear mixed-effects model revealed a significant effect of LC lesion (z = −1.96, p = 0.050). Control animals (n = 8, 104 observations) showed a mean of 0.285 ± 0.018 c-Fos positive cells per pixel, while LC lesion animals (n = 8, 105 observations) exhibited a reduction (β = −0.051, SE = 0.026) compared to controls, representing an 18% decrease relative to the control mean. Data is represented as mean ± SEM. # indicates statistical difference. **G)** Representative image of cFos immunostaining in control (WT, top and left bottom images) and LC lesion (Th-Cre, middle and right bottom images) groups. Scale bars: 500 μm (top) and 100 μm (bottom).

Taken together, our findings suggest that DA release to CA1 from LC is essential for TFC learning.

### LC lesion reduces activity and c-Fos expression of CA1 pyramidal neurons

We further investigated whether the LC lesion causes a reduction in CA1 neural activity through *in vivo* measurements of calcium sensors during TFC and immediate early gene expression induced by TFC learning.

In addition to inducing LC lesion, using the procedure described above, mice were injected with a viral vector into the CA1 to express calcium sensors in either pyramidal neurons (CaMKII-GCaMP6f) (**Fig. 2A**) or interneurons (dlx-GCaMP6f) (**Fig. 2C**). In these animals, bulk calcium activity could be measured using fiber photometry during TFC (20-s trace protocol). First, we replicated our previous findings ^13^, with control mice without LC lesion, in which an increase in the calcium signal of CA1 pyramidal neurons was observed after auditory cue presentation, during the shock and in the post-shock-1. Furthermore, lesioning the LC abolished these increases in calcium activity (**Fig. 2B**). Interestingly, the calcium activity of CA1 interneurons showed a signal enhancement profile similar to that observed in pyramidal neurons. However, lesions in the LC did not alter the activity profile of this interneuron population (**Fig. 2D**), suggesting that the LC only modulates the activity of CA1 pyramidal neurons, and not interneurons.

As a proof of concept, we measured the expression of the immediate early gene (IEG) c-Fos using immunohistochemistry to confirm the fiber photometry results (**Fig. 2E-G**). For this purpose, the brains of mice from the LC lesion group and the control group were collected 90 minutes after TFC conclusion (20-s trace protocol). After initial tissue processing, 4 to 8 slices of the dHC from each animal were stained for c-Fos. A linear mixed-effects model was used to account for repeated c-Fos measurements within subjects. The model included a random intercept for subject, with the dependent variable being cell density (cells per pixel × 100). Measurements consisted of 8–15 CA1 pyramidal layer segments per animal. The model revealed a significant effect of LC lesion, which reduced the number of c-Fos-positive cells (**Fig. 2G**).

In summary, these results suggest that LC modulates only the activity of CA1 pyramidal neurons during TFC, and not of interneurons.

### Long trace intervals produce poor learning that can be rescued by LC stimulation

To develop a robust hippocampus-dependent temporal association task in rodents, a 15-to 30-second interval between discontiguous conditioned and unconditioned stimuli is typically employed in trace fear conditioning ^18,19^. Accordingly, our previous studies have demonstrated that 20-second trace interval establishes a strong association between tone and shock in mice ^6,12,13^. Although an earlier study suggests that trace intervals should remain below 20 seconds ^7^. optimal learning within a range of trace intervals has not been systematically interrogated by other researchers under any different conditions. Thus, to examine how these parameters behave in our circumstances, we also tested the level of temporal learning in mice subjected to trace intervals of 60-, 90-, and 120-, in addition to the 20-second interval.

Therefore, in the conditioning session, independent group of mice underwent 5 trials consisting of a 20-s tone (85 dB, 3 kHz) and trace intervals of either 20-, 60-, 90-, or 120-s followed by a 2-s shock of 0.3 mA. A long 360-s intertrial interval (ITI) was established so that all animals from the different groups, especially those with the longest trace, could adequately discriminate the temporality of the trace and the ITI, thus avoiding any potential backward association between the shock and the subsequent tone of the next trial. An additional control group was exposed only to the shocks, without any tone, while maintaining the total duration of the conditioning session equivalent to that of the 20-s trace group (**Fig. 3A**). During conditioning, all groups showed a similar level of freezing in response to the tones (**Fig. S5**). The following day, associative learning was examined with 2 subsequent trials performed in a novel context by measuring the average percentage of freezing during the 20-s tone and the 20-s post-tone period corresponding to a trace-like interval. In addition, freezing levels in the first 2 minutes, before the presentation of the first tone (baseline period), was used as an indicator of contextual discrimination.

**Figure 3.**
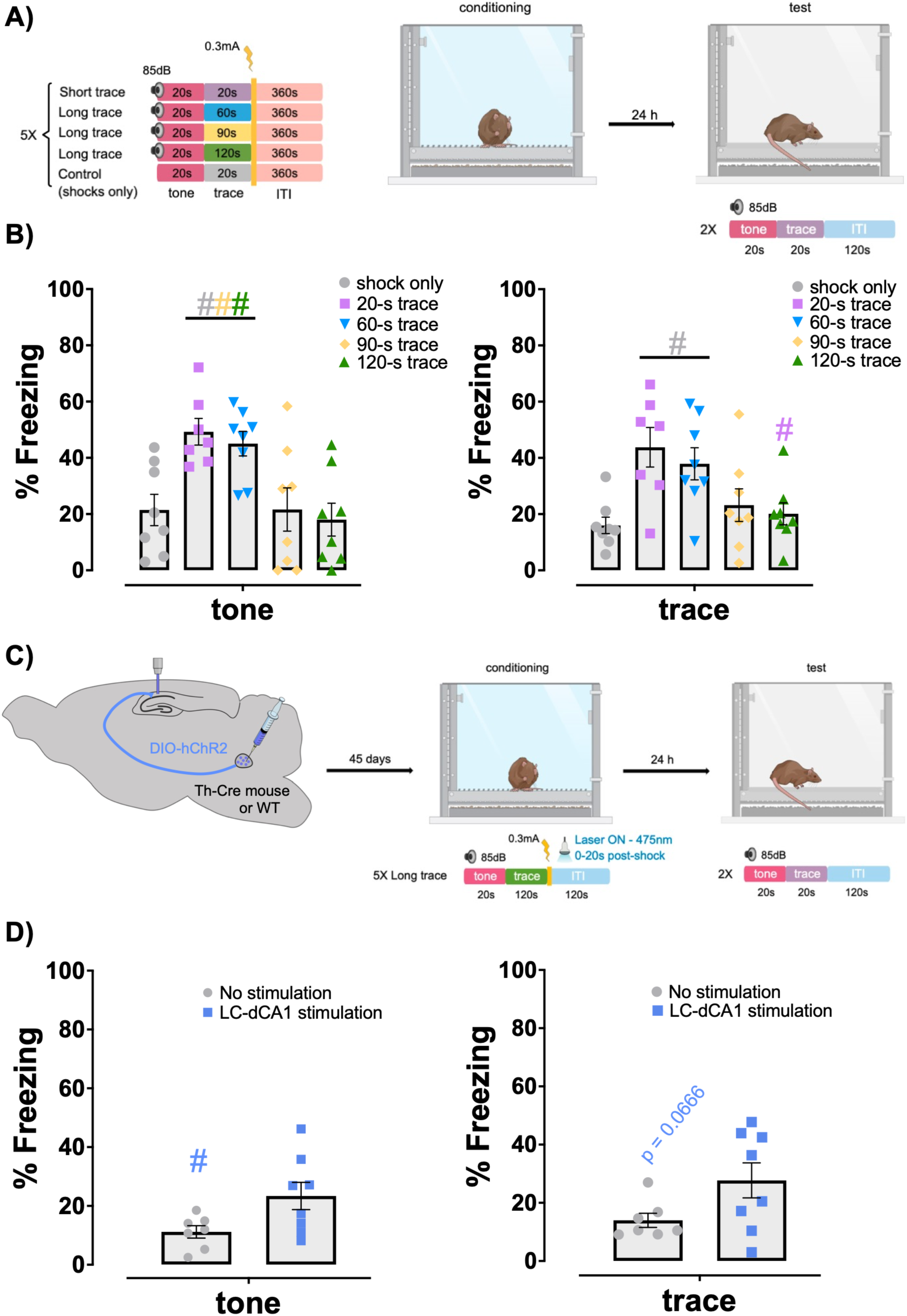
Long trace intervals produce poor learning that can be rescued by optogenetic stimulation of LC-dCA1 projections. **A)** Behavioral design illustrating distinct groups of mice that underwent variable trace intervals (20-, 60-, 90-, or 120-s) during conditioning sessions (n = 7, 8, 8, 8, respectively). Each trial was repeated five times and consisted of a 20-s tone (85 dB, 3 kHz), followed by a trace interval, a shock (0.3 mA, 2 s), and a 360-s intertrial interval (ITI). An additional group (n = 8) was exposed to the shocks without tone pairings, and its conditioning session duration was equivalent to that of the 20-s trace group. **B)** The test session was the same for all groups and consisted of 2 trials in which the tone was presented for 20-s, followed by a 20-s post-tone period corresponding to the trace-interval on the training day (trace-like) and 120-s ITI. The mean percentage of freezing during the test session was calculated for each group of animals and plotted separately for the tone and trace-like periods. One-way repeated measures ANOVA revealed a main effect of Group during the tone (F(4,34) = 6.340, p = 0.0006) and trace-like (F(4,34) = 5.199, p = 0.0022) periods. Tukey’s post-hoc test indicated that groups exposed to 20- and 60-s trace intervals differed significantly from the shock-only, 90-s trace, and 120-s trace groups during tone presentation (p < 0.05 for all comparisons). In the trace-like period, 20- and 60-s trace groups differed from the shock-only group (Tukey’s post-hoc test, p < 0.05 for both comparisons), and the 20-s trace group was different from the 120-s trace group (Tukey’s post-hoc test, p < 0.05). # indicates p < 0.05, and the color specifies the group with which the difference was significant. See also Figures S5 – S7. **C)** Cre-dependent hChR2-mCherry vector was bilaterally injected into the LC of TH-Cre (n=8) or WT (n=7) mice, and optical fibers were implanted above the dCA1. On the first day, mice underwent TFC while laser stimulation (475 nm) was delivered to CA1 (4Hz, 15ms, 10 mW) for 20 s, starting 0 s after the footshock on each training trial (5 trials in total). The next day, mice were tested in a novel context. **D)** Freezing during the test session. Th-Cre animals that had the LC-dCA1 projections stimulated during conditioning showed higher freezing levels during tone presentation (unpaired t-test, t (13) = 2.282, p = 0.0399, and a trend during the trace period (unpaired t-test, t (13) = 2.002, p = 0.0666, compared to controls, WT mice. # indicates p < 0.05. See also Figures S8 and S9. All data are expressed as mean ± SEM.

The freezing levels of the group exposed only to shocks, and not to tones, were considered a baseline measure of nonspecific non-associative learning. In order to exhibit a robust temporal association between tone and shock, the other groups would have to show higher freezing levels during tone and trace-like periods compared to the shock-only group. As expected, all groups showed low baseline freezing levels during the test session (**Fig. S6**), indicating a lack of fear generalization to the novel context. **Fig. 3B** shows that the groups exposed to trace intervals of 20- and 60-s exhibit higher freezing levels than the shock-only, 90-s, and 120-s trace groups during tone presentation. In the trace-like period, freezing levels were higher in the 20- and 60-s trace groups compared to the group subjected to shock only. Also, the 20-s trace group froze significantly more than the 120-s trace group. These results indicate that mice form stronger associative memories in our conditions when the trace interval is shorter, lasting between 20-s and 60-s. In addition, we demonstrated that doubling the number of learning trials during the conditioning session does not improve learning in the longer trace interval group (120-s) compared to shorter trace interval (20-s) (**Fig. S7**).

Given that animals in the long-trace (120-s) protocol exhibit impaired learning, we further investigated whether optogenetic stimulation of the LC-dCA1 pathway during TFC would restore associative learning. TH-Cre and WT mice were bilaterally injected with a Cre-dependent viral vector (DIO-hChR2-mCherry) into the LC to express ChR2, so that optical fibers implanted above CA1 could stimulate labeled terminals (**Fig. 3C**). Our hypothesis was that neurons encoding tone, trace, and shock had a high chance of reactivation in the post-shock period, so the optogenetic excitation of CA1-projecting LC neurons in the long-trace group would mimic the reactivation in the 20-s trace control group (**Fig. 5A-C**). For this reason, mice subjected to TFC received a 20-s laser stimulation applied to the CA1 region immediately after footshock delivery (**Fig. 3D**). During conditioning, freezing levels were not altered by optogenetic stimulation (**Fig. S8**). However, one day after TFC, the optogenetically stimulated group showed higher freezing levels during tone compared to the unstimulated group (**Fig. 3D**), although groups did not differ during the 2 min baseline (**Fig. S9**).

In addition to integrating our previous findings, these results corroborate the hypothesis that LC activation enables learning TFC by causing CA1 neurons representing different stimuli (tone, trace and shock) to be reactivated, coordinating their activity to coincide temporally despite trace gaps.

### Stimulus-specific neuronal ensembles in the dCA1 are not altered by LC lesion or longer trace intervals

Given that both DA release and calcium bulk activity depended on LC integrity, we used miniscopes to further investigate how individual CA1 pyramidal neurons behaved throughout TFC stimuli in the control (20-s trace), LC lesion, and long trace (120-s trace) groups (**Fig 4A-B**).

**Figure 4.**
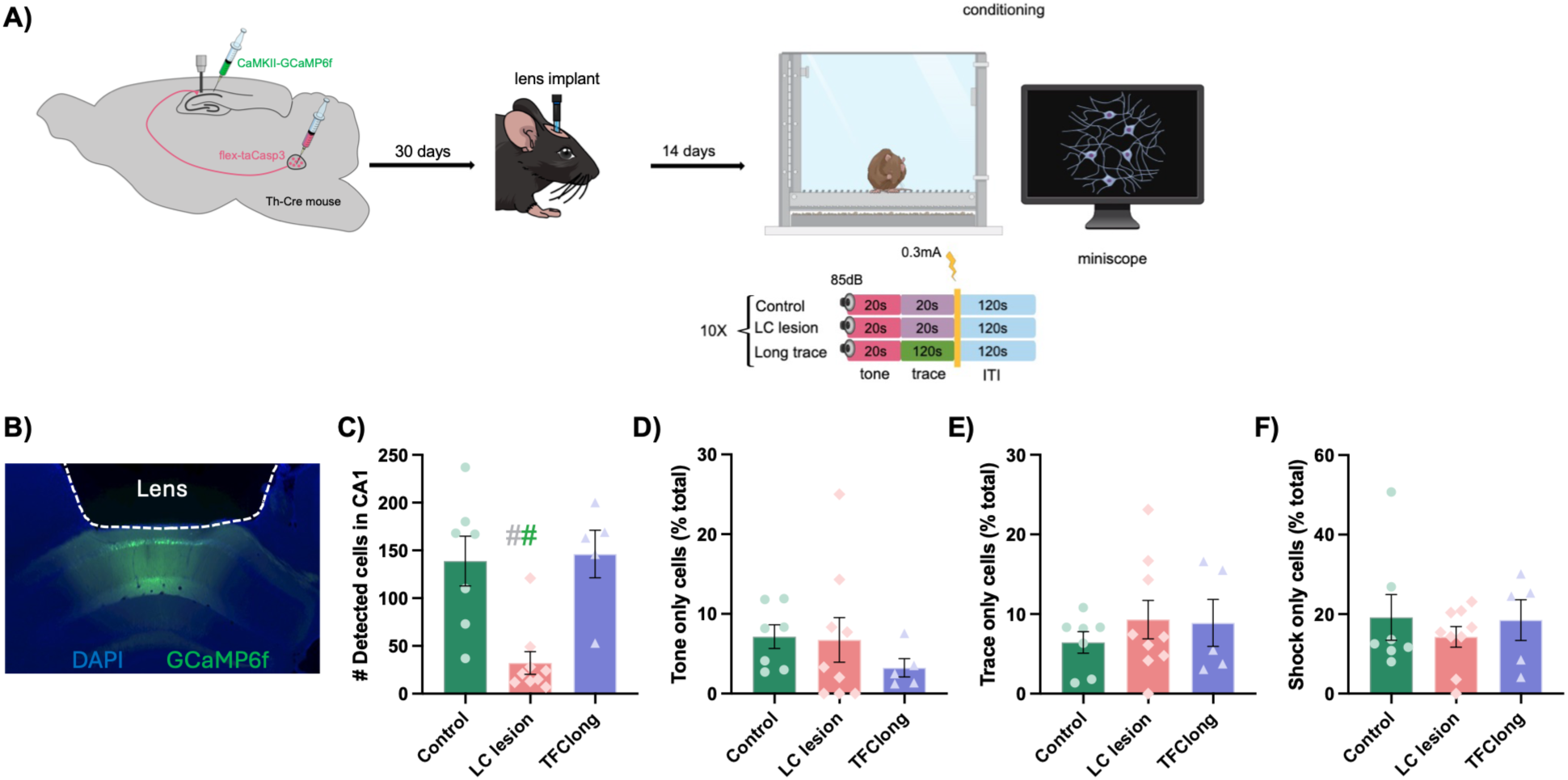
LC lesion decreases the number of detected cells in CA1, but the proportion of TFC-responsive cells remains unchanged among groups. **A)** Calcium sensor (CaMKII-GCaMP6f) was injected into dCA1. 30 days later, GRIN lenses were implanted above the CA1 region. Calcium imaging using miniscopes was performed 2 weeks later in short-trace (n = 7 animals), LC lesion (n = 9 animals) and long-trace (n = 5 animals) groups during trace fear conditioning. **B)** Representative image of GRIN lens implantation and GCaMP6f expression in the pyramidal layer of dCA1. **C)** One-way ANOVA (F (2,14) = 4.512, p = 0.0307) revealed that the number of active cells detected in CA1 was reduced in the LC lesion group (n = 290 cells in total) compared to control (n = 972 cells in total; Tukey’s post-hoc test, p = 0.00484) and long-trace groups (n = 731 cells in total; Tukey’s post-hoc test, p = 0.0482). # indicates p < 0.05. The percentage of neurons responsive to **(D)** tone-only (control: n = 81 from 972 cells [8.3%], LC lesion: 18 from 290 cells [6.2%], long-trace: 19 from 731 cells [2.6%]), **(E)** trace-only (control: n = 61 from 972 [6.3%], LC lesion: 22 from 290 cells [7.6%], long-trace: 74 from 731 cells [10.1%]), and **(F)** shock-only responsive neurons (control: n = 162 from 972 [16.7%], LC lesion: 52 from 290 cells [17.9%], long-trace: 123 from 731 cells [16.8%]) remained similar among groups (One-way ANOVA, p > 0.05). See also Figure S10. All data are expressed as mean ± SEM.

**Figure 5.**
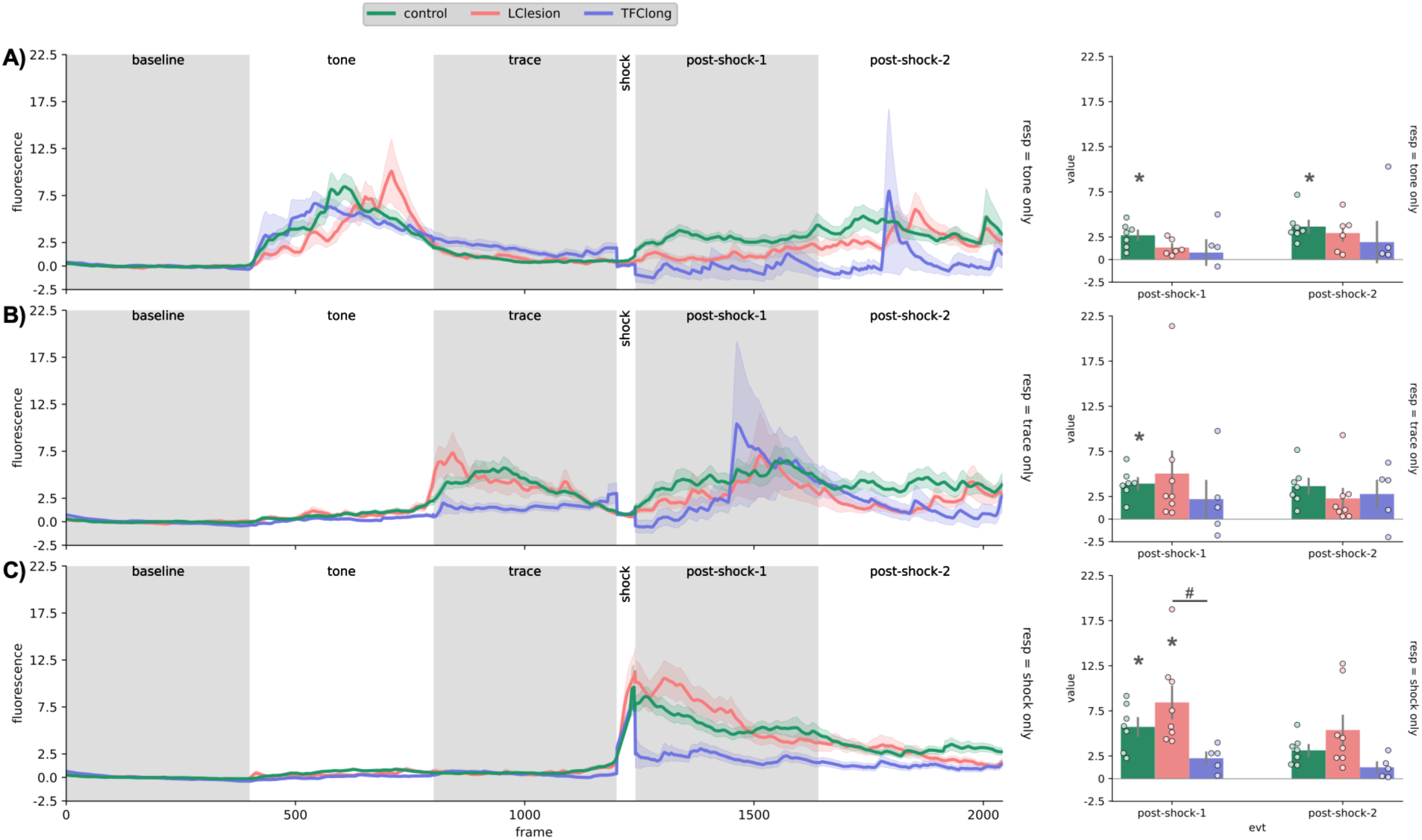
LC lesion and long-trace interval compromise post-shock neuronal reactivation. The line graphs represent the average fluorescence of all cells from control (green color), LC lesion (salmon color) and long-trace (purple color) groups during the baseline, tone, trace, shock, post-shock 1, and post-shock 2 periods. The bar graphs represent the average fluorescence (grouped by animal) during the post-shock-1 and post-shock-2 periods. **A)** Tone-only neurons from the control group (n = 81 cells from 7 animals), but not from LC lesion (n = 18 cells from 9 animals) and long-trace (n = 19 cells from 5 animals) groups, are reactivated in the post-shock-1 (t(6) = 5.2331, p = 0.0468, paired t-test with Bonferroni correction for multiple comparisons) and post-shock-2 period (t(6) = 5.7052, p = 0.0301, paired t-test with Bonferroni correction for multiple comparisons). One-way ANOVA indicated no effect of group for either post-shock-1 and post-shock-2 periods. **B)** Trace-responsive neurons from the control group (n = 61 cells from 7 animals), but not from LC lesion (n = 22 cells from 9 animals) and long-trace (n = 74 cells from 5 animals) groups, were reactivated during post-shock-1 (t(6) = 6.4704, p = 0.0155, paired t-test with Bonferroni correction for multiple comparisons). No significant effect of group was found by one-way ANOVA performed for the post-shock-1 and the post-shock-2 periods. **C)** The shock-only neurons from the control (n = 162 cells from 7 animals) and LC lesion (n = 52 cells from 9 animals) groups, but not from long-trace group (n = 123 cells from 5 animals), are reactivated in the post-shock-1 period (t(6) = 5.8982, p = 0.0253 and t (7) = 4.8168, p = 0.0462, respectively; paired t-test with Bonferroni correction for multiple comparisons). In addition, one-way ANOVA indicated a difference between groups (F (2,17) = 4.5577, p = 0.0260), with shock-only neurons from LC-lesion group showing higher calcium activity during post-shock-1 compared to long-trace group (p = 0.0203, Tukey HSD post-hoc test). # indicates p < 0.05. See also Figure S11.

In short, we first classified the cells according to the stimulus to which they responded: tone-only, trace-only, and shock-only. Additional classifications of neurons with multimodal responses (tone+trace, tone+shock, shock+trace, tone+shock+trace) are provided in supplemental information (**Fig. S10**). To do this, changes in calcium activity of each neuron was extracted across the entire conditioning session video using the Minian pipeline ^22^. The video was divided into 10 trials, each consisting of nine events: 20-s baseline, 20-s tone, 20-s trace, 2-s shock, and 5 post- shock periods of 20-s duration (post-shock-1, -2, -3, -4 and -5). The 20 seconds preceding each tone (baseline) were used for z-score normalization of the trials. Then, cells that had calcium activity two standard deviations higher in a specific epoch compared to the baseline were considered responsive to that event and classified as a subpopulation.

Our results confirmed that lesioning the LC decreased the overall number of neurons responsive to TFC (**Fig. 4C**). Interestingly, the proportion of neurons responsive only to tone (**Fig. 4D**), trace (**Fig. 4E**), and shock (**Fig. 4F**) remained unchanged. In addition, longer trace intervals did not interfere with the overall number of activated cells (**Fig. 4C**) nor with the proportion of cells responsive to specific events (**Fig. 4D-F**).

### LC lesion and longer trace intervals compromise post-shock neuronal reactivation

After classifying the cells as tone-only, trace-only, and shock-only, we sought to determine whether these neurons were reactivated throughout the post-shock phases (**Fig. 5**).

Post-shock neuronal reactivation was defined as a within-group increase in calcium activity in the post-shock periods compared to baseline. Tone-responsive neurons from the control group, but not from LC-lesion and long-trace groups, were reactivated during post-shock-1 and post-shock-2 (**Fig. 5A**). Among the trace-responsive neurons, only those from the control group were reactivated during post-shock-1 (**Fig. 5B**). In addition, neurons responsive only to shock from both the control and LC-lesion groups, but not from long-trace group, were reactivated during post-shock-1 (**Fig. 5C**). Shock-only neurons from long-trace group also showed higher calcium activity during post-shock-1 compared to LC lesion group. No differences were found between the groups for other classes of neurons during the post-shock-1 and post-shock-2 periods. Additional data showing post-shock neuronal reactivation of neurons with multimodal response are provided in the supplemental information (**Fig. S11**).

## DISCUSSION

The current study identifies the LC to dCA1 dopaminergic projection as a critical circuit for bridging temporal gaps during associative learning. We demonstrate that while trace fear conditioning (TFC) is robust at shorter intervals, associative learning decays as the temporal gap reaches 120 seconds. Our findings reveal that this learning is mediated by dopamine derived from LC, but not VTA, and facilitates the post-shock reactivation of stimulus-specific neuronal ensembles in the hippocampus. We also show that optogenetic stimulation of the LC-dCA1 pathway can rescue memory formation across prolonged temporal delays. This is similar to previous work which enhanced TFC at long trace intervals via direct optogenetic stimulation of dCA1 neurons ^7^.

### Temporal Constraints on Learning in the Hippocampus

A defining feature of the hippocampus is the ability to link events that are separated in time. While it is well-established that the dCA1 is required for this process, our data provides a clearer picture of the biological limits of this temporal bridge. We observed a significant reduction in conditioned freezing as the trace interval was extended from 20 to 120 seconds. This decay likely reflects a physiological constraint in how long a memory trace representing the conditioned stimulus (CS) can be maintained in an active (or latent) state such that it is reactivated after the unconditioned stimulus (US) occurs ^23–26^. Our results suggest that the trace interval between 90 and 120-second represents a threshold where the natural persistence of the stimulus representation is insufficient to mediate associative strengthening without additional neuromodulatory support. Similar results have been observed in vCA1 where TFC can be enhanced by increasing the release of serotonin^27^.

### Parallel to Spatial Replay

The discovery that stimulus-specific neuronal ensembles, namely tone-only and trace-only cells, reactivate immediately following a footshock provides a compelling parallel to the phenomenon of spatial replay. In the spatial domain, place cells that fire during navigation are reactivated in sequence during subsequent pauses to consolidate memory traces ^15,28,29^. Our findings suggest that the hippocampus employs a similar mechanism for temporal binding ^13^. While spatial replay links a series of locations into a coherent path, post-shock reactivation in TFC appears to link a distal auditory stimulus across a silent gap to a proximal aversive event. The fact that both LC lesions and prolonged 120-second trace intervals eliminate this reactivation indicates that temporal replay is a physiological process that links discontiguous events.

### Rehearsal, replay and memory formation

The premise that learning is a post-trial phenomenon originated with behavior theorists between the 1940s and 1960s. Tolman argued that avoidance learning might take place exclusively following an aversive event, while Leo Kamin proposed that unexpected stimuli trigger a “backward scanning” of recent memory to form associations between signals (CS) and outcomes (US) ^30,31^. This concept was later refined by Allen Wagner in the 1970s, who posited that surprising events initiate a necessary “post-trial rehearsal” process ^23–26^. Although the importance of surprise remains a pillar of modern learning theories, the assumption that it functions through post-trial rehearsal has largely faded from the field, despite historical behavioral evidence in its favor ^1,26,31–35^. Based on the work described above, we propose that surprising events enhance memory formation by inducing dopamine release at the same time the hippocampus replays recently experienced events (i.e. rehearsal) ^13^.

### Locus Coeruleus as the Primary Dopaminergic Driver in dCA1

A major point of recent debate has been the source of dopamine release in the dHC ^8–12,36^. While the VTA is the canonical source of dopamine in the brain, our selective viral-mediated lesions demonstrate that the LC is the essential driver of TFC. LC lesions significantly reduced dopamine release in the dCA1 during the tone, trace, and post-shock periods, whereas VTA lesions had a negligible impact on learning and a more limited effect on dopamine levels. These findings support the emerging view that LC-derived dopamine is the primary catecholaminergic modulator of hippocampal-dependent memory formation ^8–10,37^.

### The role of D1/D5 dopamine receptors in memory formation

Consistent with other recent findings, this study suggests that LC activity drives memory formation by releasing dopamine into the hippocampus, where it targets D1/D5 receptors ^8–10,12,37^. Activation of these receptors has been shown to strongly enhance long-term potentiation (LTP) in CA1 ^38–41^. Notably, LC stimulation does not appear to facilitate memory through hippocampal β-adrenergic receptors, a result that aligns with previous research by Thomas et al. showing these receptors are critical for the retrieval of context fear rather than initial acquisition or consolidation ^42–44^. Retrieval could be enhanced via rapid increases in intrinsic excitability, which would increase the probability that previously activated neurons (engram/memory cells) are reactivated during testing ^45^. These data suggest an interesting mechanistic divergence: DA release could facilitate the induction/maintenance of synaptic plasticity required for memory formation, while norepinephrine (NE) release enhances neuronal excitability to promote memory retrieval. This distinction would allow for the robust creation of memory representations during learning, which can then be selectively reactivated or suppressed based on their excitability state ^46,47^.

## Conclusion

In summary, these findings identify the LC-dCA1 dopaminergic circuit as an important regulator of temporal associations. By facilitating the post-shock reactivation of specific neuronal ensembles, this circuit allows the hippocampus to bridge significant gaps in time. These results not only shed light on neuromodulatory influences in hippocampus but also suggest that targeting the LC-dCA1 pathway could offer a means to enhance memory formation in conditions where temporal processing is impaired.

## STAR⋆METHODS

### KEY RESOURCE TABLE

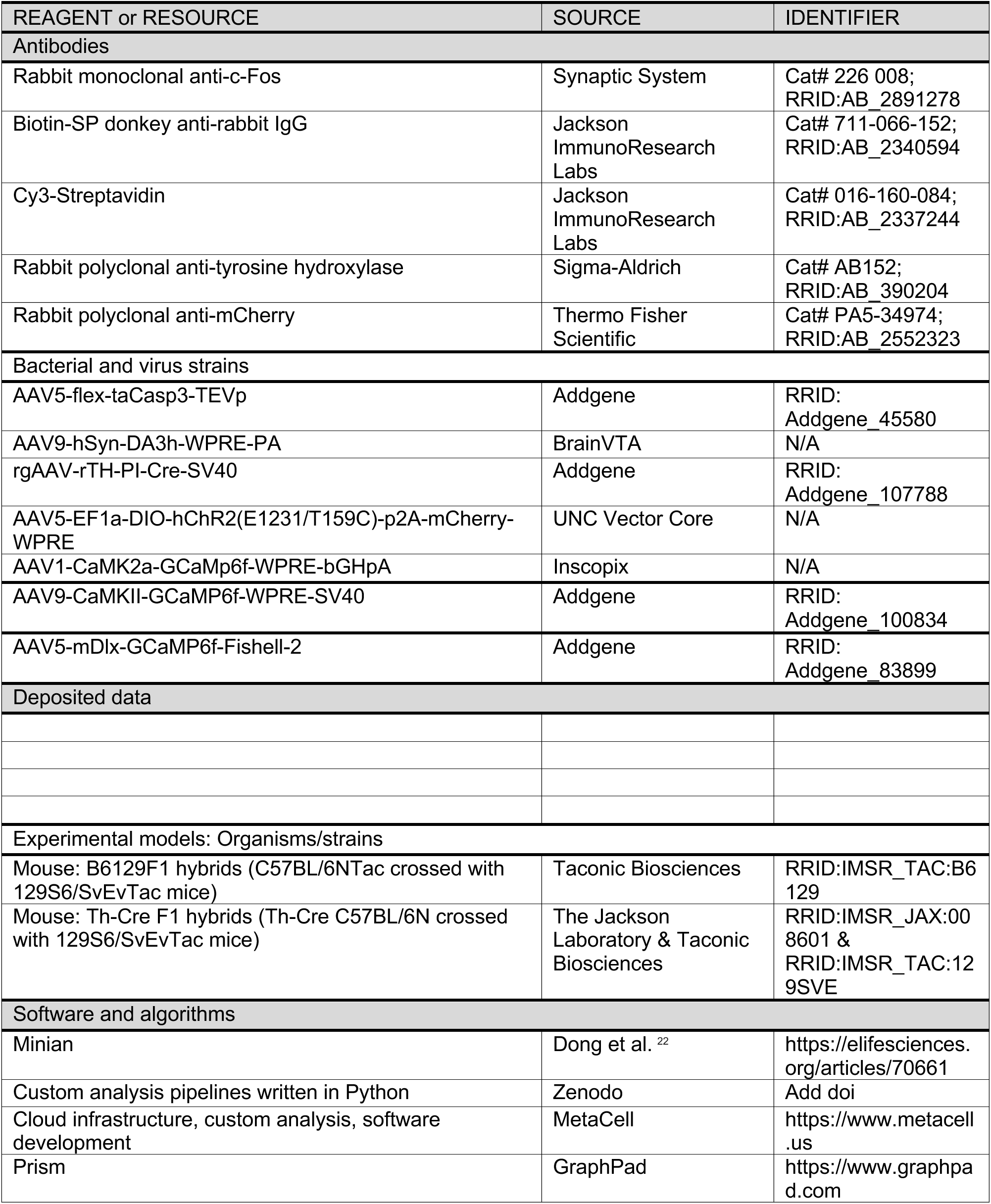
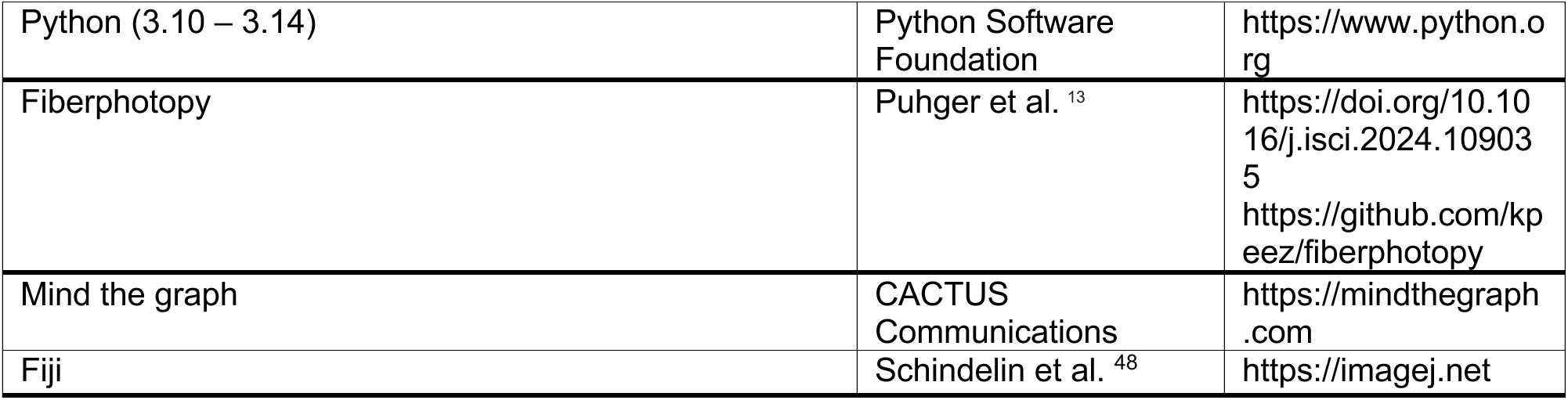

### RESOURCE AVAILABILITY

#### Lead Contact

Further information and requests for resources should be directed to Lead Contact, Brian J. Wiltgen.

#### Materials availability

This study did not generate new unique reagents.

#### Data and code availability

Requests for behavioral and/or calcium imaging data reported in this paper will be shared by the lead author upon reasonable request.

All code used for analysis of this data has been deposited at Zenodo, is publicly listed and is available as of the date of publication. The DOI is listed in the key resources table.

Any additional information required to reanalyze the data reported in this paper is available from the lead contact upon reasonable request.

### EXPERIMENTAL MODEL AND SUBJECT DETAIL

All procedures were reviewed and approved by the UC Davis Institutional Animal Care and Use Committee (IACUC; Protocols #23233, #23151, #23769). Two hybrid mouse lines were used in this study. Male B6129F1 hybrids (C57BL/6NTac crossed with 129S6/SvEvTac mice) purchased from Taconic Biosciences and male and female Th-Cre F1 hybrids (Th-Cre C57BL/6N from The Jackson Laboratory crossed with 129S6/SvEvTac mice from Taconic Biosciences) bred in the Med Neuroscience Building at the University of California, Davis. Mice were maintained on a 12 h light/12 h dark cycle with *ad libitum* access to food and water. All experiments were performed during the light portion of the light/dark cycle (0700-1900). Mice were group housed (2-4/cage) throughout the duration of the experiment. All mice were between 7 and 8 weeks old at the time of surgery and no older than 16 weeks at the endpoint of the experiments.

### METHODS DETAILS

#### Behavioral procedures

##### Trace fear conditioning

The conditioning chamber used for trace fear conditioning was housed within a sound-attenuating box (Med Associates). The chamber (30.5 cm x 24.1 cm x 21.0 cm) consisted of a front-mounted video camera, a grid floor, and overhead LED lighting capable of providing broad-spectrum and infrared light. For context A, the conditioning chamber was cleaned with 70 % and lit with both broad-spectrum and infrared light. For context B, a smooth white plastic insert was placed over the grid floor, and a curved white plastic wall was inserted into the chamber. Scented disinfectant wipes (PDI Sani-Cloth Plus) were used for cleaning, and only infrared lighting was present in the chamber, while the room lights were changed to red. A fan in the chamber and a HEPA filter in the room generated background noise (65 dB).

##### Conditioning session

In the initial set of experiments, we evaluated which trace intervals were optimal for associative learning to occur satisfactorily. In the first experiment, independent groups of animals underwent a conditioning session (context A) with 5 trials. Each trial consisted of a 20-s tone (85 dB, 3 kHz), followed by a stimulus-free trace interval of 20, 60, 90, or 120 seconds, and then a 2-s shock (0.3 mA). The intertrial interval (ITI) was of 360 seconds for all groups. In the second experiment, 5 additional trials were added to the original conditioning protocol (totaling 10), and the learning level was compared between groups with shorter (20-s) and longer (120-s) trace intervals.

Minor changes to the above protocol were adopted in subsequent experiments, as it follows: an ITI of 120-s was established in the control and LC lesion groups for fiber photometry and 1-photon (1P) calcium imaging experiments, while the 360-s trace interval was maintained in the long-trace group of the calcium imaging experiment. In these experiments, mice underwent 10 conditioning trials. In the optogenetic experiment, the same protocol from the first experiment was used (5 trials).

##### Test session

The day after the conditioning session, associative learning was examined with 2 subsequent trials performed in context B by measuring the average freezing percentage during the 20-s tone and the 20-s post-tone period corresponding to a trace-like interval. In addition, freezing levels in the first 2 minutes, before the presentation of the first tone (baseline period), were assessed to rule out fear generalization. Freezing behavior was measured using VideoFreeze software (Med Associates).

#### Surgical procedures

##### Anesthesia and analgesia

Initially, mice were deeply anesthetized with a 5 % isoflurane and oxygen mixture, and then treated with preemptive analgesia (meloxicam, 5 mg/kg s.c.). Subsequently, the mice were positioned in the digital stereotaxic and the isoflurane and oxygen mixture was maintained at a ratio of 1-2 % to sustain anesthesia. At the end of the surgery, 1 ml of 0.9 % saline s.c. was injected to prevent dehydration. Postoperative analgesia (buprenorphine, 0.1 mg/kg s.c.) was administered three times: at the end of the surgery, 24 h and 48 h later.

##### Viral microinjection and fiber implantation

Adeno-associated viruses (AAVs) were delivered at a rate of 2 nl/s via a glass pipette using a microsyringe pump (UMP3, World Precision Instruments) connected to a digital controller. After AAVs injection, the mice were kept at rest for 6 weeks (except when changes to this protocol were indicated) before the experimental behaviors were performed.

For fiber photometry experiments, 500 nl of viral vectors to express calcium (AAV9-CaMKII-GCaMP6f-WPRE-SV40 or AAV5-mDlx-GCaMP6f-Fishell-2, titer: 1×10^12^ and 1×10^13^, Addgene) or dopamine (AAV9-hSyn-DA3h-WPRE-PA, titer: 2×10^12^, BrainVTA) sensors were injected into the right CA1 (coordinates: AP –2.0 mm and ML +1.5 mm from bregma; DV –1.25 mm from dura). An optical fiber (400 μm diameter, Doric) was implanted 0.1 mm above the injection site. The fiber optic implants were secured to the skull using dental adhesive (C&B Metabond, Parkell) and dental acrylic (Bosworth Company). To lesion the brain regions of interest, a viral vector for Cre-dependent caspase-3 expression (AAV5-flex-taCasp3-TEVp, titer: 2×10^13^, Addgene) was infused into the LC (AP – 5.5 mm and ML ± 0.9 mm from bregma; DV –3.6 mm from dura) or VTA (AP −3mm and ML ± 0.6 mm from bregma; DV −4.2 mm from dura) of Th-Cre+ mice. Vehicle solution was injected into LC or VTA of Th-Cre+ mice to stablish the control non-lesioned groups. To specifically lesion the LC neurons projecting to CA1, a retrograde virus inducing Cre expression under the control of Th promoter (rgAAV-rTH-PI-Cre-SV40, titer: 1.8×10^13^, Addgene) was infused into dHC CA1, and the Cre-dependent caspase vector was infused into the LC of B6129F1 mice. The sham group was injected with the previously described retrograde virus into the CA1, while a saline-based vehicle was infused into the LC.

For 1P calcium imaging experiments, mice underwent two surgical procedures. First, the viral vector that induces expression of the calcium sensor in excitatory neurons (AAV1-CaMK2a-GCaMp6f-WPRE-bGHpA, 500 nL/hemisphere, titer: 1.3×10^12^, Inscopix) was infused into the right CA1 of B6129F1 mice. For the LC lesion group, both viral vectors to express the calcium sensor (in the CA1) and the Cre-dependent caspase-3 (in the LC) were infused into Th-Cre+ mice during the same surgical procedure. Then, 4-5 weeks later, the gradient-index (GRIN) lens was implanted above the right CA1 (see more details below).

For the optogenetic stimulation of LC neurons projecting to CA1, a viral vector for Cre-dependent ChR2 expression (AAV5-EF1a-DIO-hChR2(E123T/T159C-p2A-mCherry-WPREm titer: 4×10^12^, UNC Vector Core) was bilaterally injected (150 nl/hemisphere) into the LC and optical fibers were implanted above the dCA1. Th-Cre+ and WT mice (control) were used for this experiment.

##### Integrated gradient-index (GRIN) lens implantation

A craniotomy of 1.2 mm in diameter was performed above the right CA1 4-5 weeks after the mice were injected with viral vectors to express calcium sensors. Cortical tissue above the corpus callosum was aspirated using 27-and 30-gauge blunt needles while continuously applying chilled 0.9 % saline. Aspiration was ceased once the corpus callosum had been partially removed and bleeding had stopped. Then, the integrated GRIN lens (1 mm in diameter and 4 mm long, ProView^TM^, Inscopix) attached to the miniature microscope (Inscopix) was lowered to a depth of−1.15mm relative to the skull surface. Silicone surgical adhesive (Kwiksil, WPI) was used to fill the space between the lens and the tissue/skull. The integrated lens was fixed to the skull with dental adhesive. Mice were allowed to recover for 1-2 weeks, during which time they were house in pairs until the end of the experiment.

#### In vivo imaging of fluorescent sensors

##### Fiber photometry

Fiber photometry recordings were conducted using a Doric system, which consisted of an FPGA-based data acquisition system (Fibre Photometry Console, Doric), a programmable 4-channel LED Driver (Doric), and fluorescent Mini Cubes (Doric) in addition to a femtowatt photoreceiver (Newport 2151). Briefly, 465 nm (signal channel) and 405 nm (isosbestic channel) LEDs were respectively modulated sinusoidally at 209 and 308 Hz, with power set to ∼40 μW. Light passed through dichroic filters (Mini Cube) before transmitted into the brain via optical cables and an implanted optical fiber. The filtered light (500-540nm) emitted by the dopamine or calcium sensors was transmitted back through the optical cable and collected by the photoreceiver. Doric Neuroscience Studio software was used to modulate the LEDs and sample signals from the photoreceiver. A 12 Hz low-pass filter was applied, and the signal was decimated to 120 Hz before being written to the hard drive. The start and end of the behavioral session, as well as the times at which tones and shocks were delivered, were synchronized with the photometry acquisition system via TTL pulses controlled by the VideoFreeze software.

##### 1-photon calcium imaging through miniscopes

Neuronal calcium dynamics associated with trace fear memory at single-cell resolution were recorded in freely behaving mice using the nVista™ system (Inscopix), which consisted of the nVista miniscope camera, a commutator, and the nVista DAQ computer. Raw images were acquired at 20 frames per second during the conditioning session using Inscopix Data Acquisition Software (IDAS). Behavioral data were synchronized to the DAQ computer using TTL pulses from the VideoFreeze software that marked the beginning and end of each behavioral session, as well as the times at which tones and shocks were applied. In the control and LC lesion groups, video was recorded throughout the conditioning session. In contrast, in the long trace group, the video was stopped 120 s after each shock and restarted 20 s before the tone of the subsequent trial.

#### Immunofluorescence

##### cFos staining

90 minutes after the TFC training session, mice were transcardially perfused with PBS (6 ml/min for about 2 min), followed by 4 % PFA (6 ml/min for about 3 min). After overnight immersion in 4 % PFA, the brains were sliced using a vibratome (50 μm, coronal sections) and then stored in a freezer at −20 °C (immersed in an antifreeze solution). Firstly, slices were washed three times in PBST (phosphate-buffered saline with 0.2% Triton X-100) and incubated in donkey blocking buffer (PBST with 2% normal donkey serum) for 30min. Slices were then incubated overnight at 4 °C with a rabbit anti-c-Fos primary antibody (1:5000; Synaptic Systems, #226 008). Next, slices were incubated with biotinylated donkey anti-rabbit secondary antibody (1:500; Jackson ImmunoResearch, #711-066-152) for 1 hour, followed by streptavidin-Cy3 (1:500; Jackson ImmunoResearch, #016-160-084) for 45 minutes. DAPI (1:10000; Thermo Fisher, #D1306) counterstaining was performed for 10min. Primary and secondary antibodies were diluted with blocking buffer, while streptavidin-Cy3 and DAPI were diluted with PBS only. PBST washes were performed after antibody incubations, and PBS washes after streptavidin-Cy3 incubation and DAPI counterstaining. At last, slices were mounted on slides and cover-slipped with Vectashield anti-fade mounting media (Vector Labs, #H-1000).

##### Tyrosine hydroxylase (Th) staining

After the end of the behavior schedule (Fig. 2A), animals were perfused, and their brains were dissected and sliced as described above. The immunofluorescence procedure was the same, except that here an (rabbit) anti-Th (1:5000, Sigma, AB152) was used as the primary antibody.

##### mCherry staining

To check and better visualize hChR2-mCherry expression in projections from LC to CA1 (Fig. 6A), a rabbit anti-mCherry antibody (1:100, Thermo Fisher Scientific, PAS-34974) was used. The procedure was otherwise the same as described above.

### QUANTIFICATION AND STATISTICAL ANALYSIS

#### Immunofluorescence – image acquisition and quantification

##### cFos

Images were acquired at 20x magnification using a fluorescence slide scanner (Olympus, BX61VS). Files were opened in greyscale format and images cropped to include the pyramidal layer of the dCA1. The area of each region of interest (ROI) was measured in Fiji ^48^ and recorded. A counterbalanced subset of images was selected as the outgroup for determining the SimpylCellCounter (SCC) ^49^ parameters. Cell counting for 4-8 sections per animal (8-16 hemispheres) was performed using SCC and normalized to the ROI area. Incorrect counts were identified using the spread and average area outputs and removed from the analysis.

##### Th

Bilateral images of 2 or 3 slices of the LC or VTA were obtained for each animal, with 40x magnification, using the same slide scanner described above. So, TH staining was quantified using Fiji ^48^. Firstly, images were converted to grayscale, and then the background was adjusted before selecting the area containing TH for total pixel quantification. The bilateral measurements of each slice were added together, so that each animal sample resulted from the average of the values of all the slices.

#### Fiber photometry and 1P calcium imaging

##### Fiber photometry

All preprocessing and analysis were conducted using custom Python scripts (fiberphotopy ^13^). In brief, collected data were downsampled to 10 Hz before calculating ΔF/F. A linear regression model was fit to the isosbestic channel data (405nm) to predict the fluorescence signal of the 465nm channel. The predicted 465 nm values were then used to normalize the 465 nm signal. Recording data from individual TFC trials was isolated from the entire training session. Each trial was divided into eight periods: baseline, tone, trace, and five 20-s post-shock periods (post-shock-1, -2, -3, -4, and -5). The values were z-scored using the 20-s baseline prior to each tone. Differences in mean fluorescence values were compared within groups across periods and between groups for each period.

##### 1-photon calcium imaging through miniscopes

Minian pipeline was used to extract changes in the GCaMP6f fluorescence signal (calcium traces) across the entire raw video of conditioning session^22^. Briefly, videos were temporally downsampled to 10 frames per second by frame averaging. A Python dictionary was used to spatially downsample each video by focusing the frame on the position where cells were detected. Subsequent data processing and analysis were performed using custom-made Python codes. First, the calcium traces of the detected cells were divided into 10 trials, each consisting of 9 events: baseline (20-s pre-tone), tone, trace, and 5 20-s post-shock periods (post-shock-1, -2, - 3, -4, and -5). The 20 seconds preceding each tone (baseline) were used for z-score normalization of the trials. Then, cells were sorted based on their ability to respond to a specific stimulus or to multimodal stimuli. Cells that had activity two standard deviations higher to a specific event compared to the baseline were considered responsive to that event and classified as one of the following subpopulations: tone-only, trace-only, shock-only, tone-trace, tone-shock, trace-shock and tone-trace-shock.

In further analysis, cell reactivation was defined as the condition in which a responsive subpopulation of cells exhibited statistically higher activity in the post-shock period than in the baseline.

#### Statistical analyses

Sample sizes were based on previous studies using similar methods. Prior to experimentation, animals were randomly assigned to groups. Normally distributed data was analyzed using paired or unpaired t-test (two-sided), one or two-way repeated-measures ANOVA followed by Tukey or Bonferroni post hoc test corrected for multiple comparisons as appropriate. Linear mixed-effect models were used to account for repeated immunofluorescence measurements (Th, cFos and mCherry staining) in the same animal.

For the behavioral data, freezing percentage was measured during each trial period (baseline, tone, trace, post-shocks) and averaged across trials for each animal. Differences in group performance in each trial period were compared independently.

For fiber photometry and 1P calcium imaging experiments, sequential time periods (baseline, tone, trace, post-shocks) from the conditioning session were treated as repeated measures.

A threshold of p < 0.05 was used to determine statistical significance. Statistical analyses were performed in Python or GraphPad Prism version 10, and all figures were generated in Python, GraphPad Prism or Mind the Graph.

## Acknowledgments

We thank MetaCell, especially Phil Dong, for assisting in customizing the pipelines for miniscope data analysis. We also would like to thank Nipunika Bandara and Noah Michelson for their assistance in cell counting in the c-Fos experiment.

## Author contributions

Conceptualization, validation, methodology and data curation: A.P.C., C.R.A.F.D. and BJW; formal analysis and visualization: A.P.C. and C.R.A.F.D.; investigation: A.P.C., C.R.A.F.D., D.C. and S.M.; implementation of the computer code and supporting algorithms: A.P.C. and MetaCell; writing original draft: A.P.C. and BJW; review & editing: all authors; project administration: A.P.C. and BJW; funding acquisition, resources and supervision: BJW.

## Declaration of interest

All authors declare no competing interests.

## Funding

This work was supported by grants #R01NS129217 from NINDS to BJW.

## SUPPLEMENTAL INFORMATION

Document S1. Figures S1–S11.

## Notes

### Competing Interest Statement

The authors have declared no competing interest.

## References

1. Kamin, L.J. (1961). Trace conditioning of the conditioned emotional response. J. Comp. Physiol. Psychol. 54, 149–153. 10.1037/h0045611.

2. McEchron, M.D., Bouwmeester, H., Tseng, W., Weiss, C., and Disterhoft, J.F. (1999). Hippocampectomy disrupts auditory trace fear conditioning and contextual fear conditioning in the rat. Hippocampus 8, 638–646. 10.1002/(SICI)1098-1063(1998)8:6<638::AID-HIPO6>3.0.CO;2-Q.

3. Chowdhury, N., Quinn, J.J., and Fanselow, M.S. (2005). Dorsal hippocampus involvement in trace fear conditioning with long, but not short, trace intervals in mice. Behavioral Neuroscience 119, 1396–1402. 10.1037/0735-7044.119.5.1396.

4. Raybuck, J.D., and Lattal, K.M. (2014). Bridging the interval: Theory and neurobiology of trace conditioning. Behavioural Processes 101, 103–111. 10.1016/j.beproc.2013.08.016.

5. Kitamura, T., Macdonald, C.J., and Tonegawa, S. (2015). Entorhinal–hippocampal neuronal circuits bridge temporally discontiguous events. Learning & Memory 22, 438– 443. 10.1101/lm.038687.115.

6. Wilmot, J.H., Puhger, K., and Wiltgen, B.J. (2019). Acute Disruption of the Dorsal Hippocampus Impairs the Encoding and Retrieval of Trace Fear Memories. Front. Behav. Neurosci. 13. 10.3389/fnbeh.2019.00116.

7. Sellami, A., Al Abed, A.S., Brayda-Bruno, L., Etchamendy, N., Valério, S., Oulé, M., Pantaléon, L., Lamothe, V., Potier, M., Bernard, K., et al. (2017). Temporal binding function of dorsal CA1 is critical for declarative memory formation. Proceedings of the National Academy of Sciences 114, 10262–10267. 10.1073/pnas.1619657114.

8. Kempadoo, K.A., Mosharov, E. V., Choi, S.J., Sulzer, D., and Kandel, E.R. (2016). Dopamine release from the locus coeruleus to the dorsal hippocampus promotes spatial learning and memory. Proceedings of the National Academy of Sciences 113, 14835– 14840. 10.1073/pnas.1616515114.

9. Wagatsuma, A., Okuyama, T., Sun, C., Smith, L.M., Abe, K., and Tonegawa, S. (2018). Locus coeruleus input to hippocampal CA3 drives single-trial learning of a novel context. Proceedings of the National Academy of Sciences 115. 10.1073/pnas.1714082115.

10. Takeuchi, T., Duszkiewicz, A.J., Sonneborn, A., Spooner, P.A., Yamasaki, M., Watanabe, M., Smith, C.C., Fernández, G., Deisseroth, K., Greene, R.W., et al. (2016). Locus coeruleus and dopaminergic consolidation of everyday memory. Nature 537, 357–362. 10.1038/nature19325.

11. McNamara, C.G., and Dupret, D. (2017). Two sources of dopamine for the hippocampus. Trends Neurosci. 40, 383–384. 10.1016/j.tins.2017.05.005.

12. Wilmot, J.H., Diniz, C.R., Crestani, A.P., Puhger, K.R., Roshgadol, J., Tian, L., and Wiltgen, B.J. (2024). Phasic locus coeruleus activity enhances trace fear conditioning by increasing dopamine release in the hippocampus. Elife 12. 10.7554/eLife.91465.3.

13. Puhger, K., Crestani, A.P., Diniz, C.R.A.F., and Wiltgen, B.J. (2024). The hippocampus contributes to retroactive stimulus associations during trace fear conditioning. iScience 27, 109035. 10.1016/j.isci.2024.109035.

14. McNamara, C.G., Tejero-Cantero, Á., Trouche, S., Campo-Urriza, N., and Dupret, D. (2014). Dopaminergic neurons promote hippocampal reactivation and spatial memory persistence. Nat. Neurosci. 17, 1658–1660. 10.1038/nn.3843.

15. Ambrose, R.E., Pfeiffer, B.E., and Foster, D.J. (2016). Reverse Replay of Hippocampal Place Cells Is Uniquely Modulated by Changing Reward. Neuron 91, 1124–1136. 10.1016/j.neuron.2016.07.047.

16. Singer, A.C., and Frank, L.M. (2009). Rewarded Outcomes Enhance Reactivation of Experience in the Hippocampus. Neuron 64, 910–921. 10.1016/j.neuron.2009.11.016.

17. Dupret, D., O’Neill, J., Pleydell-Bouverie, B., and Csicsvari, J. (2010). The reorganization and reactivation of hippocampal maps predict spatial memory performance. Nat. Neurosci. 13, 995–1002. 10.1038/nn.2599.

18. Huerta, P.T., Sun, L.D., Wilson, M.A., and Tonegawa, S. (2000). Formation of Temporal Memory Requires NMDA Receptors within CA1 Pyramidal Neurons. Neuron 25, 473– 480. 10.1016/S0896-6273(00)80909-5.

19. McEchron, M.D., Bouwmeester, H., Tseng, W., Weiss, C., and Disterhoft, J.F. (1999). Hippocampectomy disrupts auditory trace fear conditioning and contextual fear conditioning in the rat. Hippocampus 8, 638–646. 10.1002/(SICI)1098-1063(1998)8:6<638::AID-HIPO6>3.0.CO;2-Q.

20. Raybuck, J.D., and Lattal, K.M. (2014). Bridging the interval: Theory and neurobiology of trace conditioning. Behavioural Processes 101, 103–111. 10.1016/j.beproc.2013.08.016.

21. Quinn, J.J., Oommen, S.S., Morrison, G.E., and Fanselow, M.S. (2002). Post-training excitotoxic lesions of the dorsal hippocampus attenuate forward trace, backward trace, and delay fear conditioning in a temporally specific manner. Hippocampus 12, 495–504. 10.1002/hipo.10029.

22. Dong, Z., Mau, W., Feng, Y., Pennington, Z.T., Chen, L., Zaki, Y., Rajan, K., Shuman, T., Aharoni, D., and Cai, D.J. (2022). Minian, an open-source miniscope analysis pipeline. Elife 11. 10.7554/eLife.70661.

23. Wagner, A.R. Evolution of a structured connectionist model of Pavlovian conditioning (AESOP). In Cognitive Processes in Animal Behavior (Routledge).

24. Wagner, A.R. (2016). Priming in STM: An Information-Processing Mechanism for Self-Generated or Retrieval-Generated Depression in Performance. In Habituation (Routledge), pp. 95–128. 10.4324/9781315636726-3.

25. Wagner, A.R. (2018). Expectancies and the priming of STM. Cognitive Processes in Animal Behavior, 177–209. 10.4324/9780203710029-7/EXPECTANCIES-PRIMING-STM-ALLAN-WAGNER.

26. Wagner, A.R., Rudy, J.W., and Whitlow, J.W. (1973). Rehearsal in animal conditioning. J. Exp. Psychol. 97, 407–426. 10.1037/h0034136.

27. Zhao, Y., Li, X., Chang, T., Xie, S., Wu, Y., Li, S., Cao, J., Wang, D., Deng, F., Feng, J., et al. (2025). Serotonin shapes the temporal window for associative fear learning. Preprint, 10.1101/2025.08.06.669013 https://doi.org/10.1101/2025.08.06.669013.

28. Mallory, C.S., Widloski, J., and Foster, D.J. (2025). The time course and organization of hippocampal replay. Science (1979). 387, 541–548. 10.1126/science.ads4760.

29. Kleinman, M.R., and Foster, D.J. (2025). Spatial localization of hippocampal replay requires dopamine signaling. Elife 13. 10.7554/ELIFE.99678.2.

30. Tolman, E.C. (1948). Cognitive maps in rats and men. Psychol. Rev. 55, 189–208. 10.1037/h0061626.

31. Kamin, L.J. (1969). Predictability, Surprise, Attention, and Conditioning. In Punishment Aversive Behavior, pp. 279–296.

32. Rescorla, R.A. (1968). Probability of shock in the presence and absence of cs in fear conditioning. J. Comp. Physiol. Psychol. 66, 1–5. 10.1037/h0025984.

33. Wagner, A.R., and Terry, W.S. (1975). Backward conditioning to a CS following an expected vs. a surprising UCS. Anim. Learn. Behav. 3, 370–374. 10.3758/BF03213461.

34. Terry, W.S., and Wagner, A.R. (1975). Short-term memory for “surprising” versus “expected” unconditioned stimuli in Pavlovian conditioning. J. Exp. Psychol. Anim. Behav. Process. 1, 122–133. 10.1037/0097-7403.1.2.122.

35. Joos, E., Vansteenwegen, D., and Hermans, D. (2012). Post-acquisition repetitive thought in fear conditioning: An experimental investigation of the effect of CS-US-rehearsal. J. Behav. Ther. Exp. Psychiatry 43, 737–744. 10.1016/j.jbtep.2011.10.011.

36. Broussard, J.I., Yang, K., Levine, A.T., Tsetsenis, T., Jenson, D., Cao, F., Garcia, I., Arenkiel, B.R., Zhou, F.-M., De Biasi, M., et al. (2016). Dopamine Regulates Aversive Contextual Learning and Associated In Vivo Synaptic Plasticity in the Hippocampus. Cell Rep. 14, 1930–1939. 10.1016/j.celrep.2016.01.070.

37. Gálvez-Márquez, D.K., Salgado-Ménez, M., Moreno-Castilla, P., Rodríguez-Durán, L., Escobar, M.L., Tecuapetla, F., and Bermudez-Rattoni, F. (2022). Spatial contextual recognition memory updating is modulated by dopamine release in the dorsal hippocampus from the locus coeruleus. Proceedings of the National Academy of Sciences 119. 10.1073/pnas.2208254119.

38. Huang, Y.Y., and Kandel, E.R. (1995). D1/D5 receptor agonists induce a protein synthesis-dependent late potentiation in the CA1 region of the hippocampus. Proc. Natl. Acad. Sci. U. S. A. 92, 2446–2450. 10.1073/pnas.92.7.2446.

39. Duszkiewicz, A.J., McNamara, C.G., Takeuchi, T., and Genzel, L. (2019). Novelty and Dopaminergic Modulation of Memory Persistence: A Tale of Two Systems. Trends Neurosci. 42, 102–114. 10.1016/j.tins.2018.10.002.

40. Otmakhova, N.A., and Lisman, J.E. (1996). D1/D5 dopamine receptor activation increases the magnitude of early long-term potentiation at CA1 hippocampal synapses. J. Neurosci. 16, 7478–7486. 10.1523/JNEUROSCI.16-23-07478.1996.

41. Fuchsberger, T., Stockwell, I., Woods, M., Brzosko, Z., Greger, I.H., and Paulsen, O. (2025). Dopamine increases protein synthesis in hippocampal neurons enabling dopamine-dependent LTP. Elife 13. 10.7554/eLife.100822.3.

42. Murchison, C.F., Zhang, X.-Y., Zhang, W.-P., Ouyang, M., Lee, A., and Thomas, S.A. (2004). A Distinct Role for Norepinephrine in Memory Retrieval. Cell 117, 131–143. 10.1016/S0092-8674(04)00259-4.

43. Zhang, L., Ouyang, M., Ganellin, C.R., and Thomas, S.A. (2013). The Slow Afterhyperpolarization: A Target of β 1 -Adrenergic Signaling in Hippocampus-Dependent Memory Retrieval. The Journal of Neuroscience 33, 5006–5016. 10.1523/JNEUROSCI.3834-12.2013.

44. Ouyang, M., and Thomas, S.A. (2005). A requirement for memory retrieval during and after long-term extinction learning. Proceedings of the National Academy of Sciences 102, 9347–9352. 10.1073/pnas.0502315102.

45. Thomas, S.A. (2015). Neuromodulatory signaling in hippocampus-dependent memory retrieval. Hippocampus 25, 415–431. 10.1002/hipo.22394.

46. Pignatelli, M., Ryan, T.J., Roy, D.S., Lovett, C., Smith, L.M., Muralidhar, S., and Tonegawa, S. (2019). Engram Cell Excitability State Determines the Efficacy of Memory Retrieval. Neuron 101, 274–284.e5. 10.1016/j.neuron.2018.11.029.

47. Frankland, P.W., Josselyn, S.A., and Köhler, S. (2019). The neurobiological foundation of memory retrieval. Nat. Neurosci. 22, 1576–1585. 10.1038/s41593-019-0493-1.

48. Schindelin, J., Arganda-Carreras, I., Frise, E., Kaynig, V., Longair, M., Pietzsch, T., Preibisch, S., Rueden, C., Saalfeld, S., Schmid, B., et al. (2012). Fiji: an open-source platform for biological-image analysis. Nat. Methods 9, 676–682. 10.1038/nmeth.2019.

49. Bal, A., Maureira, F., and Arguello, A.A. (2020). SimpylCellCounter: an automated solution for quantifying cells in brain tissue. Sci. Rep. 10, 12570. 10.1038/s41598-020-68138-4.

