## Supplemental Information for "Dopamine Release from the Locus Coeruleus Enables Temporal Binding in the Hippocampus"

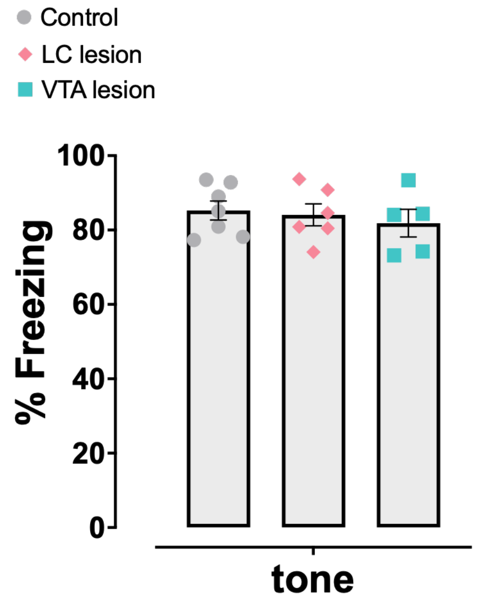


**Figure S1 –** Groups did not differ in freezing levels during the conditioning session (One-way ANOVA, $p > 0.05$; N = 7, 6 and 5).

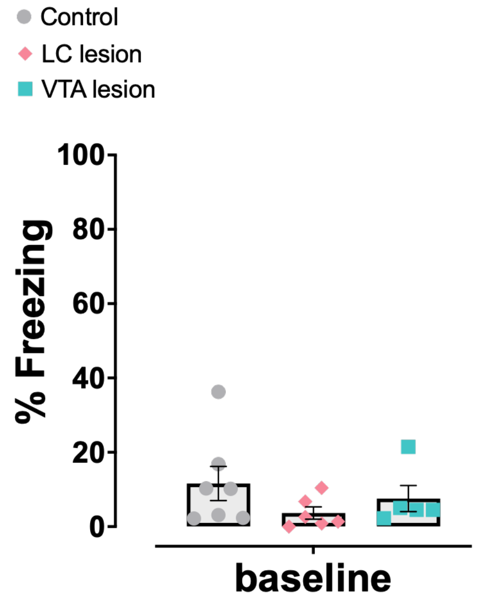


**Figure S2 –** Groups did not differ in freezing levels during the 2 min baseline period of the test session (One-way ANOVA, $p > 0.05$; N = 7, 6 and 5).

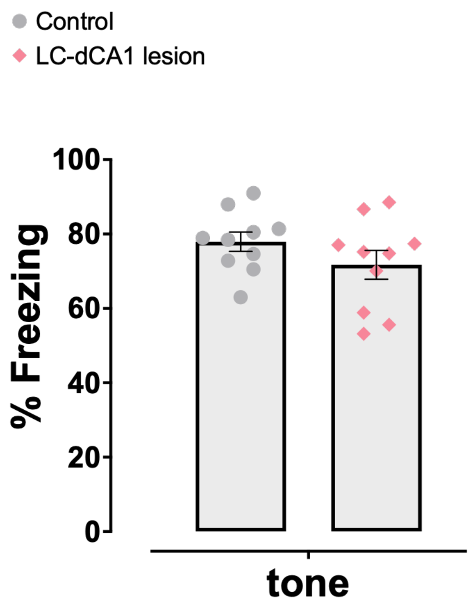


**Figure S3 –** Freezing levels did not differ between groups during the conditioning session (unpaired t-test, $p > 0.05$; N = 10 and 10).

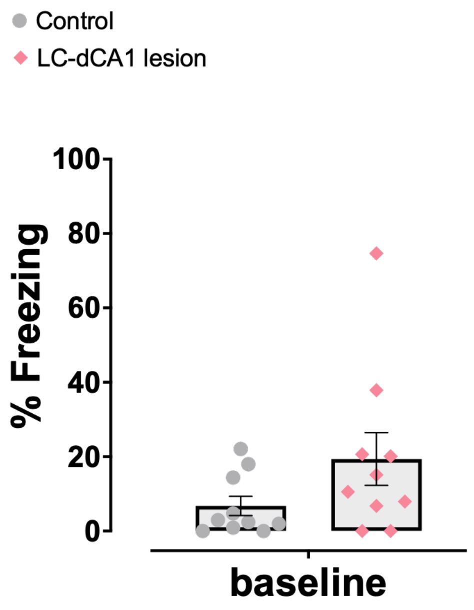


**Figure S4 –** Freezing levels did not differ between groups during the 2 min baseline of the test session (unpaired t-test, $p > 0.05$; N = 10 and 10).

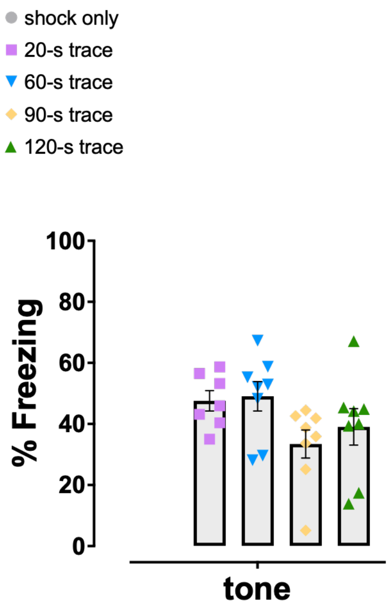


**Figure S5 –** During conditioning, all groups showed a similar level of freezing in response to the tones (One-way ANOVA, $p > 0.05$; N = 7, 8, 8, 8).

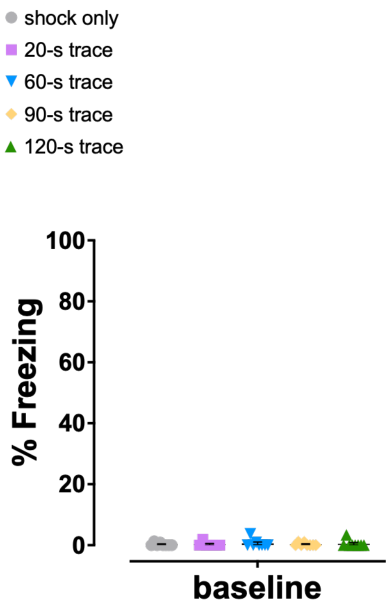


**Figure S6 –** All groups showed low baseline freezing levels during the test session (context B), indicating a lack of fear generalization to the novel context (One-way ANOVA, $p > 0.05$; N = 7, 8, 8, 8).

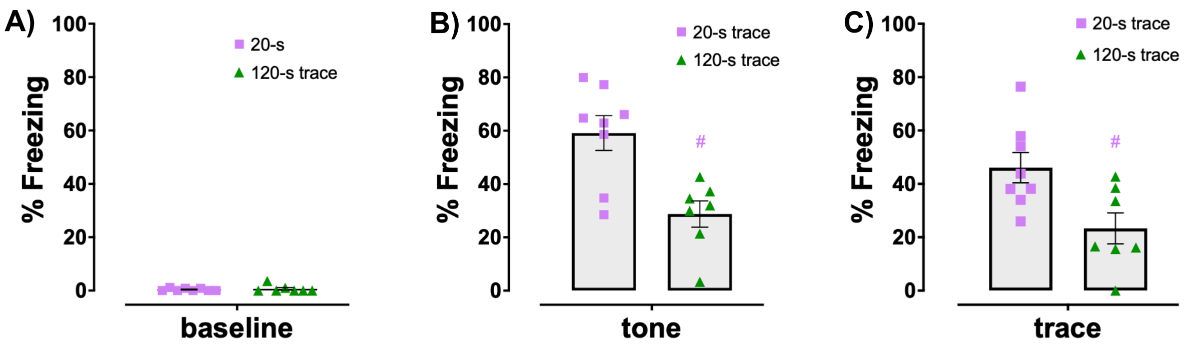


**Figure S7 – Increasing the number of trials during conditioning session does not compensate the impairment caused by the long trace interval.**

**A)** Freezing levels did not differ between groups during the baseline period of the test session (unpaired t-test, $p > 0.05$; N = 8 and 7). Unpaired t-test indicated that 120-s trace group froze less than 20-s trace group during tone ($t(13) = 3.627$, $p = 0.0031$, **B**) and trace periods ($t(13) = 2.791$, $p = 0.0153$, **C**).

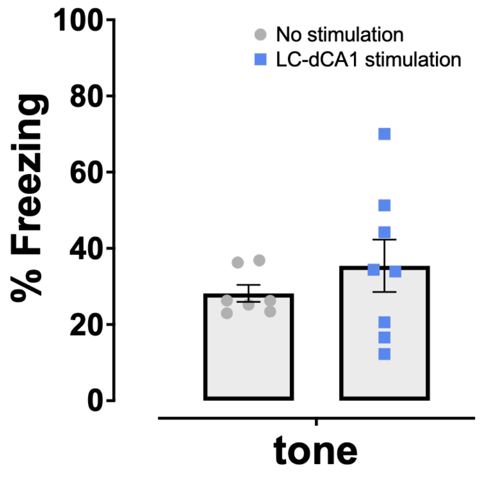


**Figure S8 –** Freezing levels did not differ between groups during the conditioning session (unpaired t-test, $p > 0.05$). N = 7 and 8, following the order in which the data were presented.

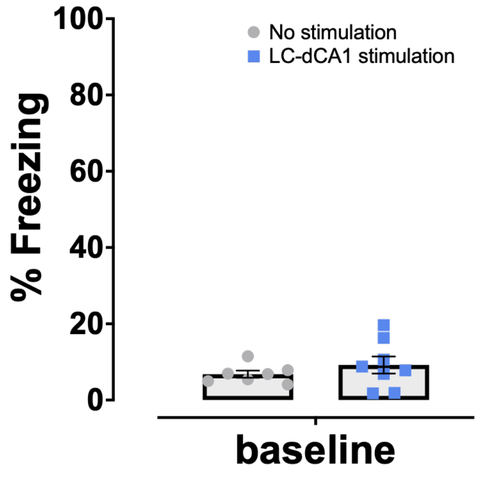


**Figure S9 –** Freezing levels did not differ between groups during the 2 min baseline of the test session (unpaired t-test, $p > 0.05$). N = 7 and 8, following the order in which the data were presented.

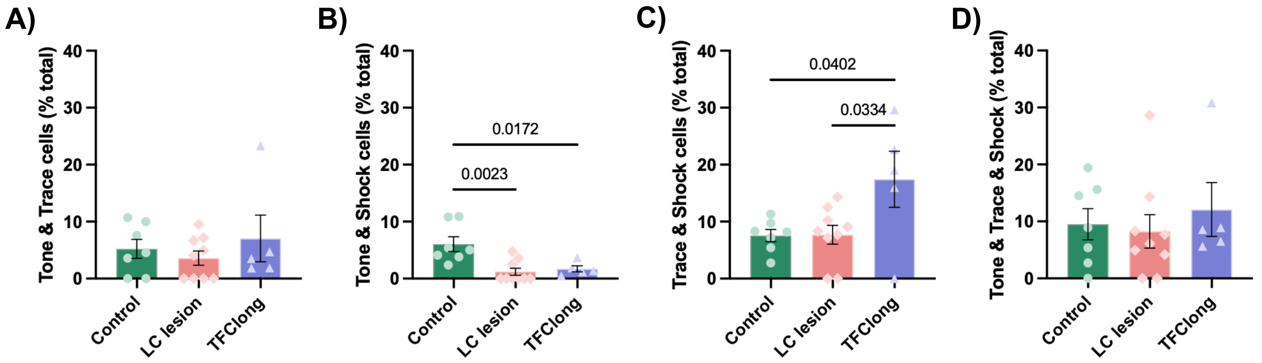


**Figure S10 – Classification of neurons with multimodal response.**

The percentage of **(A)** tone+trace (control: n = 65 from 972 cells [6.7%], LC lesion: 13 from 290 cells [4.5%], long-trace: 21 from 731 cells [2.9%]) , **(C)** tone+trace+shock neurons (control: n = 113 from 972 cells [11.6%], LC lesion: 17 from 290 cells [5.9%], long-trace: 95 from 731 cells [13.0%]) remained similar among groups (One-way ANOVA, $p > 0.05$). **(B)** For the tone+shock neurons, one-way ANOVA indicated a difference between groups ($F (2,18) = 8.8480$, $p = 0.0021$), with LC lesion (n = 5 from 290 cells, 1.7% of group total) and long-trace (n = 10 from 731 cells, 1.4% of group total) groups showing a lower percentage of neurons compared to control group (n = 50 from 972 cells, 5.1% of group total) ($p = 0.0023$ and $p = 0.0172$, respectively; Tukey post-hoc test). **(D)** For the shock+trace neurons, one-way ANOVA indicated a difference between groups ($F (2,18) = 4.5520$, $p = 0.0251$), with long-trace group (n = 159 from 731, 21.8% of group total) showing a higher percentage of neurons compared to control (n = 82 from 972 cells, 8.4% of group total) and LC lesion group (n = 26 from 731 cells, 9.0% of group total) ($p = 0.0402$ and $p = 0.0334$, respectively; Tukey post-hoc test).

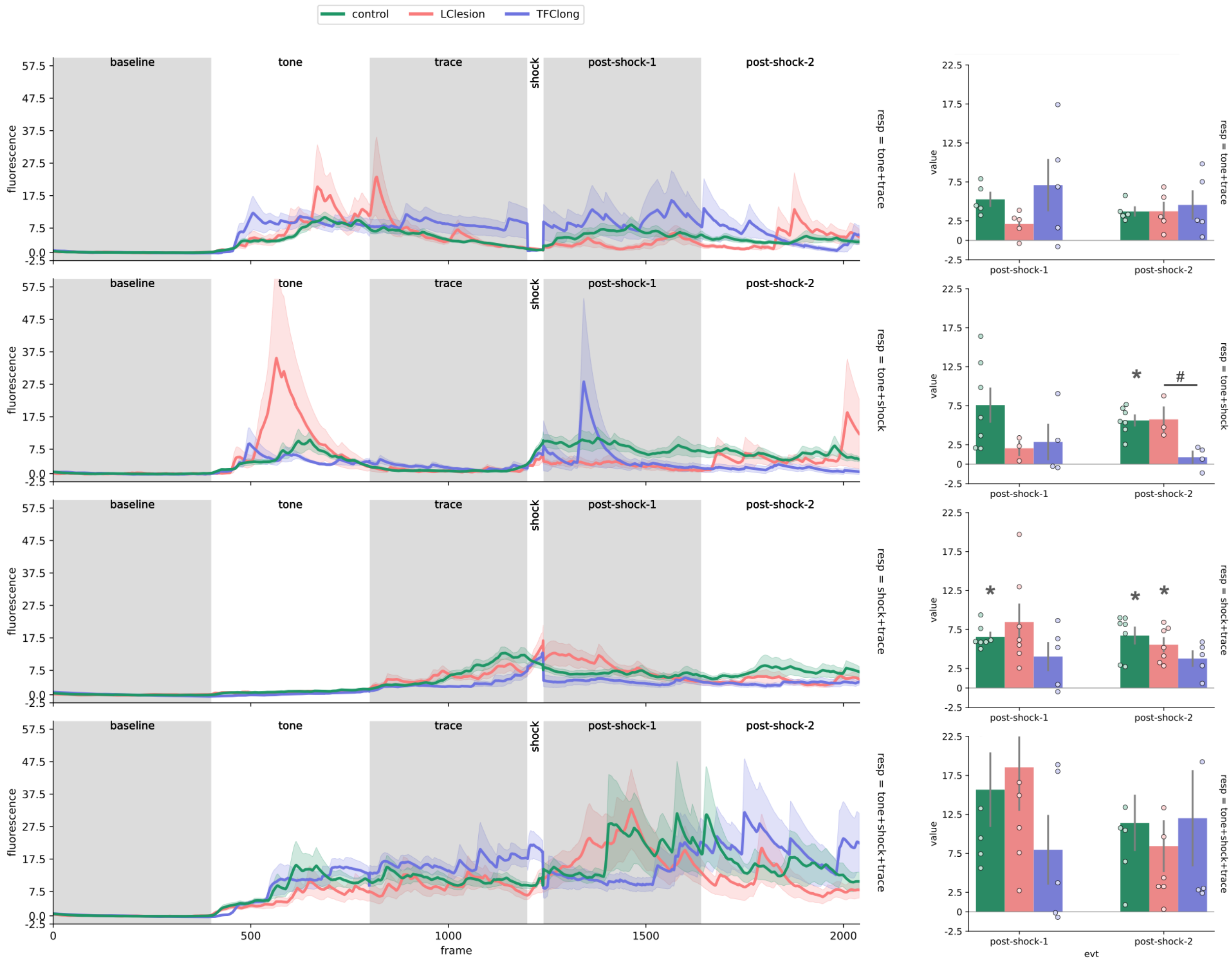


**Figure S11 – Post-shock neuronal reactivation of multimodal neurons.**

The line graphs represent the average fluorescence of all cells from control (green color), LC lesion (salmon color) and long-trace (purple color) groups during the baseline, tone, trace, shock, post-shock 1, and post-shock 2 periods. The bar graphs represent the average fluorescence (grouped by animal) during the post-shock-1 and post-shock-2 periods.

**A)** Tone+trace neurons from all groups (n = control: 65 cells from 7 animals, LC lesion: 13 cells from 9 animals and long-trace: 21 cells from 5 animals), are not reactivated in the post-shock-1 and post-shock-2 period. One-way ANOVA indicated no group effect for either post-shock-1 and post-shock-2 periods.

**B)** Tone+shock responsive neurons from the control group (n = 50 cells from 7 animals), but not from the LC lesion (n = 5 cells from 9 animals) and long-trace (n = 10 cells from 5 animals) groups, were reactivated during post-shock-2 ($t(6) = 8.4561$, $p = 0.0036$, paired t-test with Bonferroni correction for multiple comparisons). In addition, one-way ANOVA indicated a difference between groups ($F(2,11) = 9.1366$, $p = 0.0046$), with LC-lesion group showing higher calcium activity compared to long-trace group ($p = 0.0156$, Tukey HSD post-hoc test).

**C)** Shock+trace neurons from the control (n = 82 cells from 7 animals), but not from the LC lesion (n = 26 cells from 9 animals) and long-trace (n = 159 cells from 5 animals) groups, are reactivated in the post-shock-1 period ($t(6) = 11.8531$, $p = 0.0005$). In addition, shock+trace neurons from control and LC lesion groups are reactivated in the post-shock-2 period ($t(6) = 6.4855$, $p = 0.0153$ and $t(7) = 6.4798$, $p = 0.0154$, respectively; paired t-test with Bonferroni correction for multiple comparisons). One-way ANOVA indicated no effect of group for either post-shock-1 and post-shock-2 periods.

**D)** Tone+shock+trace neurons from all groups (n = control: 113 cells from 7 animals, LC lesion: 17 cells from 9 animals and LC lesion: 95 cells from 5 animals) are not reactivated in the post-shock-1 and post-shock-2 periods. One-way ANOVA indicated no effect of group for either post-shock-1 and post-shock-2 periods.
